# Mathematical AI Predicts 40% and 57% Influenza Vaccine Effectiveness Against H1N1 and H3N2, Respectively, for the 2026-–2027 U.S. Season

**DOI:** 10.64898/2026.10.22.760980

**Authors:** Xiang Xiang Wang, Guo-Wei Wei

**Author notes:** Corresponding author: Guo-Wei Wei.

## Abstract

**Background:** Clinical influenza vaccine effectiveness (VE) has often been low over the past two decades, highlighting the need for preseason VE forecasting. Timely forecasts could help health authorities plan prevention strategies and allocate healthcare resources. However, preseason forecasting of clinical influenza VE remains relatively uncommon.

**Methods:** We obtained hemagglutinin (HA) sequences from GISAID for U.S. specimens collected in August–September before each influenza season from 2010–2011 through 2024–2025. After quality screening, we compared each virus with the corresponding vaccine target strain for the same season and subtype or lineage using two mathematical distances based on amino acid patterns in the HA protein. We calculated mean distances for each season and subtype or lineage and paired them with published clinical VE estimates, yielding 26 observations across 13 seasons. Using a mathematical AI approach, we fitted linear regression models of VE against each distance and evaluated their predictive performance by forecasting later seasons using data from earlier seasons, with the historical mean VE for each subtype or lineage as a reference. We then applied the same approach to 87 HA sequences collected in August–September 2026 to forecast VE for the 2026–2027 season.

**Results:** We generated component-specific forecasts of influenza VE for the upcoming 2026–2027 U.S. season. Both mathematical distances were negatively associated with historical VE, indicating that greater differences between circulating viruses and vaccine strains were associated with lower VE. The topological model forecast VE of 40.3%, 56.9%, and 47.1% against A(H1N1)pdm09, A(H3N2), and B/Victoria, respectively, whereas the geometric model forecast corresponding VE values of 34.0%, 56.8%, and 51.1%.

## 1 Introduction

Influenza vaccination is an important means of reducing seasonal illness, but its effectiveness varies from one season to another [1]. Knowing the likely level of protection before the influenza season would help public health authorities prepare for a potentially substantial influenza burden, plan healthcare capacity, refine risk communication, and inform public health decision-making, particularly when protection against a specific subtype is expected to be limited. Clinical vaccine effectiveness (VE) estimates, however, require vaccination records and laboratory test results from people who develop respiratory illness and seek medical care [2]. These data accumulate as the season progresses. Interim estimates can be reported during the season: the 2025–2026 US interim report was published on 12 March 2026 using surveillance data collected from September 2025 through February 2026 [3]. Full-season journal reports may appear substantially later. For example, reports for the 2017–2018, 2023–2024 and 2024–2025 seasons were first published online on 2 February 2019, 6 January 2025 and 14 July 2026, respectively [4–6]. These dates refer to journal publication rather than the first public availability of any VE information; the Centers for Disease Control and Prevention (CDC) also presented preliminary end-of-season estimates for 2025–2026 in June 2026 [7]. These estimates measure protection using observed clinical outcomes rather than forecast protection before those outcomes occur. Earlier forecasts could complement clinical surveillance by providing additional time to prepare for lower protection, with the potential to reduce infections and avoidable healthcare costs.

Virus sequences offer an information source that can be examined before sufficient clinical outcomes have accumulated. Influenza viruses undergo genetic and antigenic evolution [8], and changes at specific HA positions can alter antigenic properties [9]. Differences between circulating and vaccine viruses are relevant to protection: the 2025–2026 US interim report documented the predominance of antigenically drifted A(H3N2) viruses alongside lower interim VE than in recent seasons [3]. Hemagglutinin (HA) sequences describe variation in a major surface protein. Experimental studies have linked changes at specific HA positions to antigenic changes [9], and sequence-based antigenic distances have been related to historical vaccine protection [10]. However, larger sequence differences do not necessarily imply larger antigenic differences [8]. VE also depends on factors such as prior immunity, vaccine characteristics, the study population and the clinical outcome assessed [5, 6]. A key challenge is determining whether distances between preseason HA sequences and vaccine strains can provide useful information for predicting VE.

Mathematical artificial intelligence (AI), particularly topological deep learning [11], offers an effective approach for biomolecular analysis [12]. Mathematical representations of biomolecular data of this kind are reviewed by Nguyen et al. [13], and persistent topological Laplacians have been applied to the structural analysis of viral proteins [14]. Topological deep learning and spectral representations have also been used to study how viral mutations affect viral infectivity and antibody resistance [15–17]. These approaches lead to the successful predictions of crucial spike protein mutation sites [15] and emerging SARS-CoV-2 dominant variants [18, 19]. More recently, Hozumi and Wei introduced *k*-mer topology to compare sequences through persistent topological and spectral features, and examined its use for antigenic-distance analysis of SARS-CoV-2 variants [20].

In this work, we propose the use of mathematical distances between vaccine target and influenza variants to predict influenza VE. Our study combines HA sequences from GISAID [21] with published clinical VE estimates to forecast VE for a future season. We use topological and geometric methods to quantify differences between preseason HA sequences and vaccine strains, then relate these distances to historical clinical VE through regression. We construct two distances from the positions of short amino-acid fragments, or *k*-mers, along HA. The topological distance compares how occurrences of each fragment form connected groups across scales [20]. The geometric distance represents topological and Laplacian spectral descriptors as subspaces on a Grassmann manifold and compares these subspaces using chordal distance [22, 23]. Grassmann-manifold representations of this kind have also been used for other high-dimensional biological data [24], and related persistent-Laplacian constructions extend to sheaf-theoretic settings [25]. These constructions summarize sequence variation in different ways. Evaluating them on the same sequences, vaccine targets and clinical observations allows us to compare their relationships with VE.

Using August–September sequences to forecast next season VE requires addressing two questions. First, do virus–vaccine target distances measured before the season reflect those observed during the subsequent clinical enrollment period? Second, can the relationship between sequence distance and historical VE help predict effectiveness in a later season? These questions are related but distinct. Agreement between collection windows supports the timing of the sequence measurement, whereas prediction against clinical outcomes assesses its value for forecasting protection. Examining both provides a basis for evaluating preseason sequence distances as early indicators of VE.

We pair US HA sequences from specimens collected during August and September with published component-specific VE estimates of the next season, yielding 26 season–component observations from 13 historical influenza seasons between 2010–2011 and 2024–2025. After screening sequences and verifying the clinical estimates, we calculate distances to the corresponding vaccine strains and fit a separate regression for each distance. We assess prediction on later seasons using pooled models and distance normalisation fitted to earlier data. We also compare topological distances calculated from August–September sequences with those from sequences collected during the subsequent VE-study enrollment window, to assess whether preseason measurements reflect later-season virus–vaccine differences. We then forecast VE against A(H1N1)pdm09, A(H3N2) and B/Victoria for the 2026– 2027 US influenza season using 87 eligible HA sequences from specimens collected in the United States during August–September 2026. These forecasts are intended for comparison with final US Flu VE Network estimates for medically attended, laboratory-confirmed outpatient influenza, matched by subtype or lineage. This comparison target differs from the aggregate overall VE on CDC’s seasonal summary webpage.

## 2 Mathematical foundations

We describe each HA sequence in two steps. First, the positions at which short fragments recur define a family of graphs: increasing the allowed gap joins separate occurrences into connected groups. Second, the resulting measurements are compared either as vectors or as linear subspaces. The first view retains differences in their numerical values; the second compares the subspaces they span. We first introduce the topological objects, then construct them from *k*-mer positions, before describing the geometry of subspaces and the two distance functions.

### 2.1 Basic topological concepts

Topology describes features such as connectivity and holes. For finite data, simplicial complexes provide a way to describe these features using simple building blocks [26, Chapter III]. A vertex is a 0-simplex, an edge is a 1-simplex, a filled triangle is a 2-simplex, and a solid tetrahedron is a 3-simplex (Figure 1a). These standard examples also introduce the basic objects in the persistent-Laplacian treatment of Wei and Wei [27].

**Figure 1:**
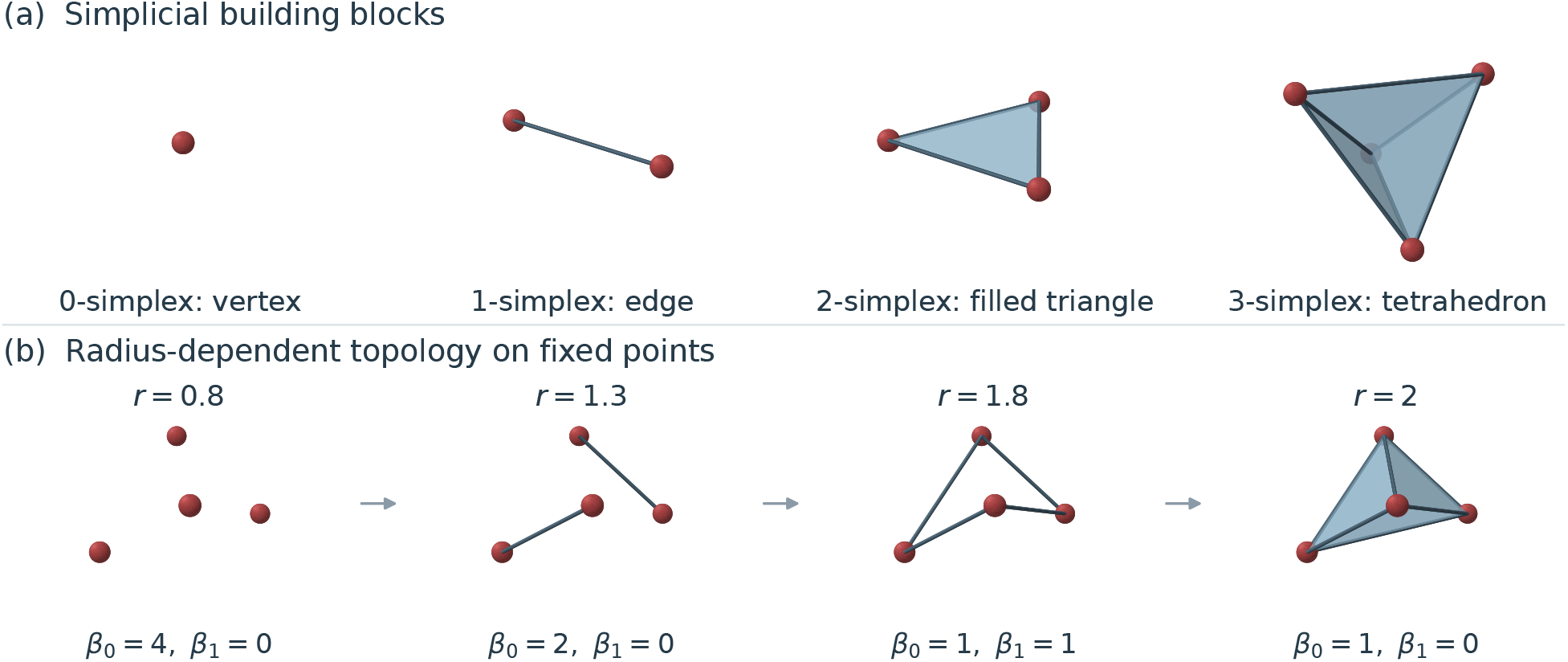
Simplices and a radius-based filtration. (a) Basic simplices of dimensions zero to three. (b) Vietoris–Rips complexes on the same four non-coplanar points, shown from the same three-dimensional viewpoint. An edge enters when its Euclidean length is at most *r*, and every clique defines a simplex. Increasing *r* joins four isolated points into two components and then one loop; adding faces fills the loop, and the final complex is a solid tetrahedron. The displayed Betti numbers count components and independent loops. Shaded faces show the three-dimensional structure; sphere and edge thicknesses are used only for visibility. All panels schematically illustrate basic topological concepts.

More generally, a *q*-simplex is the convex hull of *q* + 1 affinely independent vertices *v*_0_, …, *v*_*q*_:

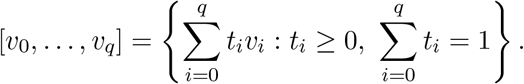

A face is the simplex obtained from a subset of these vertices. A simplicial complex is a collection of simplices containing all their faces, with any nonempty intersection shared as a common face. Its 1-skeleton consists of its vertices and edges and is a graph. Vertices belong to the same connected component when an edge path joins them.

Homology describes connectivity and higher-dimensional holes algebraically [26, Chapter IV]. Over a field ?, the *q*th Betti number is *β*_*q*_ = dim *H*_*q*_(*K*; ?) for a complex *K*. In low dimensions, *β*_0_ counts connected components, *β*_1_ counts independent loops and *β*_2_ counts enclosed cavities. An unfilled triangular boundary has one loop; adding its triangular face fills that loop. A simplex itself has one connected component and no higher-dimensional holes. These distinctions explain why vertices, edges and filled faces must be specified separately.

The scale at which points are connected determines which of these features are visible. To examine more than one scale, let *X* = *{x*_1_, …, *x*_*n*_*}* ⊂ ℝ^*d*^ be a finite point cloud. Its Vietoris–Rips complex connects points according to a distance threshold [26, Section III.2]. Here, *r* denotes the edge-length threshold, so the complex contains precisely the vertex sets whose pairwise distances are at most *r*:

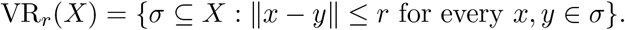

Thus a close pair supplies an edge, three mutually close points supply a filled triangle, and four mutually close points supply a 3-simplex. This rule specifies the faces as well as the edges; drawing an edge cycle alone is not the same as filling its interior.

For *r*_0_ ≤ *r*_1_ ≤ · · · ≤ *r*_*m*_, every simplex present at one radius remains present at larger radii, giving a filtration [26, Chapter VII]

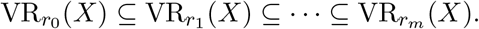

The point coordinates remain fixed throughout; it is their connection structure that changes. Figure 1b illustrates this process on four points. At *r* = 0.8 they form four separate components. At *r* = 1.3, two edges leave two components. At *r* = 1.8, a connected cycle has formed, so *β*_0_ = 1 and *β*_1_ = 1. At *r* = 2, the additional edges and faces fill the loop and complete a tetrahedron, leaving *β*_0_ = 1 and *β*_1_ = 0. This is the radius-dependent construction illustrated in the persistent-Laplacian examples of Wei and Wei [27].

Persistent homology tracks the appearance and disappearance of homological features along this filtration [26, Chapter VII]. A feature is born when it first appears and dies when it merges with an older component or becomes a boundary. For example, adding edges can merge components, whereas adding triangular faces can remove loops. The interval between birth and death records persistence across connection scales, rather than elapsed biological time. Recording only *β*_*q*_(*r*) gives a Betti curve: a count of the features present at each radius, without retaining their individual identities.

These examples introduce the general construction. In the present study, the points are occurrence positions along an HA sequence, and the distances use component counts and graph-Laplacian measurements across radii. The loops and filled faces above explain the broader topological language; higher-dimensional Betti numbers are not inputs to the VE models. The next subsection specifies the one-dimensional position sets and their graphs.

### 2.2 k-mer positions and their topology

Following the position-based representation of Hozumi and Wei [20], let *A* be a finite alphabet with *A* = |*A*|, and let *S* = *S*_1_*S*_2_ · · · *S*_*L*_ ∈ *A*^*L*^ be a sequence. For an integer *k* ≥ 1 the set of *k*-mer types is *A*^*k*^, of cardinality *A*^*k*^. For a *k*-mer type *l* ∈ *A*^*k*^ define its occurrence set

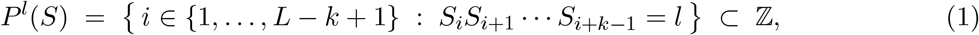

a finite point cloud on the integer line recording where in *S* that *k*-mer occurs. A window containing a character outside *A* — a nucleotide ambiguity code, or a non-standard residue — matches no type and so belongs to no occurrence set. The indexed collection 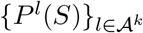 records the positions of each *k*-mer type without requiring an alignment between sequences.

This construction applies to any finite alphabet. Here, *A* is the set of 20 standard amino acids; an individual amino acid, an adjacent pair and an adjacent triplet correspond to *k* = 1, 2, 3. The source methods provide the position-based construction and its nucleotide implementations [20, 23]; the protein settings used here are specified in Section 3.2.

For a radius *r* ≥ 0, join two distinct positions *i, j* ∈ *P* ^*l*^(*S*) when |*i* − *j*| ≤ *r*. This gives a graph 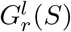the 1-skeleton of the Vietoris–Rips complex at scale *r*. Positions are compared along the sequence as written; Hozumi and Wei [20] also describe a circular variant that joins the two ends, which is not used here. Every set of pairwise connected vertices specifies a simplex, but the vertices and edges alone determine the component count. Increasing *r* produces the filtration described above: 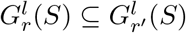whenever *r* ≤ *r*^*′*^.

The zeroth homology space 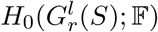, over a field ?, has one independent class for each component. Its dimension is therefore the component count. Equivalently, let 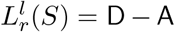A be the graph Laplacian, where A is the adjacency matrix and D contains the vertex degrees on its diagonal [28]. The multiplicity of the zero eigenvalue equals the number of connected components, giving

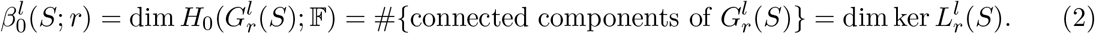

The function 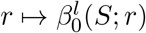is the Betti-0 curve. For a nonempty occurrence set it decreases from the number of occurrences to one. For positions *{*1, 3, 8*}*, the gap of two joins the first two points, and the gap of five joins the remaining point. The curve thus retains information about separation as well as occurrence count.

Because the point cloud lies in one dimension, this curve has a closed form.

#### Proposition 1

(Betti-0 of a one-dimensional point cloud). Let *p*_1_ *< p*_2_ *<* · · · *< p*_*n*_ be the elements of *P* ^*l*^(*S*) in increasing order and let *g*_*i*_ = *p*_*i*+1_ − *p*_*i*_ for 1 ≤ *i* ≤ *n* − 1 be the consecutive gaps. Then for every *r* ≥ 0,

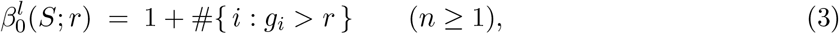

and 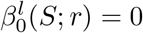when *n* = 0.

The position-based filtration is that of Hozumi and Wei [20]; in one dimension each consecutive gap larger than *r* separates two components, which gives the closed form above.

Proposition 1 allows the Betti-0 curve to be evaluated without constructing a simplicial complex or a graph Laplacian. After sorting the positions and their consecutive gaps in *O*(*n* log *n*) time, evaluation at *T* radii costs *O*(*T* log *n*) by binary search on the sorted gaps.

The Betti-0 curve reads only the harmonic part of the spectrum — the multiplicity of the eigenvalue 0. The non-harmonic spectrum, consisting of the positive eigenvalues [22, 27], carries complementary information about how tightly each cluster of occurrences is connected, not merely how many clusters there are. Write 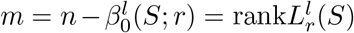for the number of strictly positive eigenvalues and, when *m* ≥ 1, define the first two moments of the non-harmonic spectrum

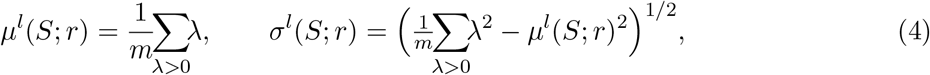

the sums running over the positive eigenvalues of 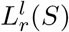with multiplicity. These too avoid an eigendecomposition.

#### Proposition 2

(Degree-only form of the spectral moments). Let deg_*i*_ be the degree of vertex *i* in 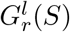. Then

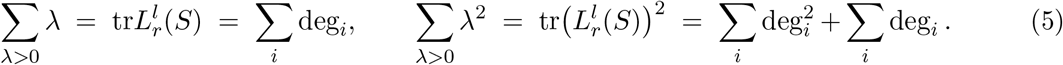

Consequently *µ*^*l*^ and *σ*^*l*^ are determined by the degree sequence alone and, after sorting, are computable in *O*(*n*) per radius.

These identities follow from the standard trace relations for the combinatorial graph Laplacian [28].

The smallest positive eigenvalue 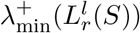 supplies a further spectral summary [22, 28]. When several nontrivial components are present, it is the minimum of their smallest positive eigenvalues. Implementation details are given in Supplementary Information, Section C.4.

Fix a finite grid of radii *r*_1_ *< r*_2_ *<* · · · *< r*_*T*_ and a set of channels *G*, each channel being one of the scalar summaries above:

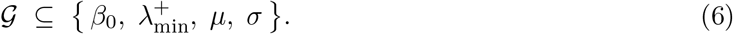

For a sequence *S*, a channel *g* ∈ *G* and an order *k*, define the feature block

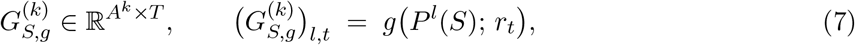

one row per *k*-mer type in lexicographic order and one column per radius. The collection 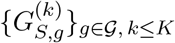is the common object from which both predictors are derived. Each matrix has a word-type axis and a radius axis. This common arrangement lets us compare its entries directly or compare the subspaces obtained after combining the measurements.

The blocks are sparse: when *A*^*k*^ ≫ *L*, a 566-residue protein contains at most 564 distinct 3-mers out of 20^3^ = 8,000 possible types, so most rows vanish identically. Rows that vanish for every sequence being compared may be omitted from sparse computations, provided their common coordinates and the row-count convention used in normalisation are preserved. The entries of different channels and different *k* have incommensurable units and magnitudes, so the blocks must be normalised before they are combined; both distances below do so, and both therefore require a scale estimated over a corpus of comparisons rather than over a single pair.

### 2.3 Subspaces and Grassmannian geometry

A nonzero vector determines a line through the origin; two independent vectors determine a plane. Changing the lengths or the basis vectors need not change that line or plane. Grassmannian geometry treats the subspace itself as the object, separating it from the particular vectors used to represent it. We use the standard definitions below [29, 30].

#### Definition 1

(Grassmann manifold). For 1 ≤ *ρ < D*, the real Grassmann manifold is [29]

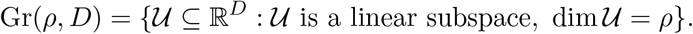

A point of this manifold is an entire *ρ*-dimensional subspace, rather than a point lying in that subspace.

Thus Gr(1, 3) contains the unoriented lines through the origin in ℝ ^3^, while Gr(2, 3) contains the planes through the origin. More generally, Gr(*ρ, D*) is a compact smooth manifold of dimension *ρ*(*D* − *ρ*) [29].

To perform calculations, choose an orthonormal basis for the subspace and place its vectors in the columns of *U* . Such matrices form the Stiefel manifold [29]

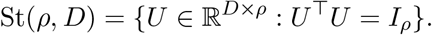

The columns are mutually perpendicular and have unit length, but this representation is not unique. Matrices *U* and *V* span the same subspace exactly when *V* = *UO* for an orthogonal matrix *O* ∈ *O*(*ρ*) = *{O* ∈ ℝ^*ρ×ρ*^ : *O*^⊤^*O* = *I*_*ρ*_*}*. Consequently,

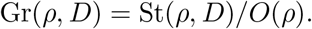

The quotient identifies all orthonormal bases of a common subspace [29]. Rotating an orthonormal basis within its plane does not move the Grassmann point.

The orthogonal projector gives a basis-independent representation [31, Section 2.5]:

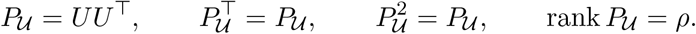

For any *x* ∈ ℝ ^*D*^, *P*_*U*_ *x* is the closest point in *U* to *x*. Because (*UO*)(*UO*)^⊤^ = *UU*^⊤^, every choice of orthonormal basis produces the same projector.

Two subspaces of the same dimension can also be compared through their principal angles [31, Section 6.4.3]. For orthonormal bases *U, V*, let *η*_1_ ≥ · · · ≥ *η*_*ρ*_ be the singular values of *U*^⊤^*V* . The principal angles are

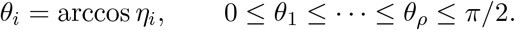

For two lines this becomes the acute angle *θ* = arccos |*u*^⊤^*v*| between unit representatives; reversing either vector does not change it. For higher-dimensional subspaces, successive angle pairs are chosen orthogonally to the preceding pairs within each subspace. All angles are zero when the subspaces coincide. These projectors and angles provide two equivalent ways to express the chordal distance below.

### 2.4 Betti-curve Euclidean distance

The first distance compares sequences directly through their Betti-0 curves, using only the channel *G* = {*β*_0_}. At each radius, a curve records how many groups of occurrences remain disconnected for a particular *k*-mer type. At radius zero this is its occurrence count; as the radius increases, the curve also reflects the gaps separating those occurrences. For example, positions *{*1, 3, 8*}* have consecutive gaps 2 and 5, so their Betti-0 curve equals 3 for 0 ≤ *r <* 2, 2 for 2 ≤ *r <* 5, and 1 for *r* ≥ 5. Comparing these curves therefore incorporates both occurrence counts and spatial separation along the sequence.

For a fixed order *k*, the block 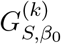 contains one row per *k*-mer type and one column per sampled radius. The operation vec arranges all entries into a single vector, with the same ordering used for every sequence. The within-order distance is

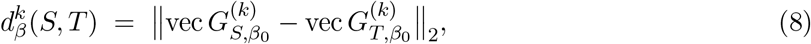

where the Euclidean norm takes the square root of the sum of squared entrywise differences. Equivalently, this is the Frobenius norm of the difference between the two feature blocks [31, Sections 2.2–2.3]. Each matched *k*-mer–radius coordinate contributes to the comparison; larger discrepancies contribute more strongly through squaring.

Different orders capture patterns at different word lengths, and their raw distances need not have comparable magnitudes. We therefore combine them as

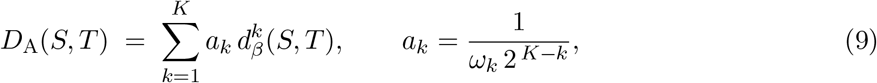

where *ω*_*k*_ *>* 0 is the mean of 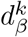 over a specified corpus of comparisons. Division by *ω*_*k*_ expresses each distance relative to its reference magnitude. The additional factor 2^−(*K*−*k*)^ assigns progressively greater weight to higher orders: it doubles when *k* increases by one. These two factors have distinct roles, normalising the distance scale and setting the relative emphasis across orders. This construction follows the *k*-mer topological distance of Hozumi and Wei [20] in their Betti-only setting, with the spectral term switched off. Their own antigenic-distance application retains a spectral channel alongside the Betti curves, which is one reason we also evaluate a distance built from the nonzero spectrum. The scale-fitting populations used in this study are specified in the methods.

Because the weights are positive and each within-order comparison is a Euclidean distance, *D*_A_ is nonnegative and symmetric. It does not separate sequences: distinct sequences can have identical sampled Betti curves, so a zero distance means equality of the chosen topological features rather than of the sequences themselves. In particular, the curves summarise connectivity rather than retaining the absolute locations of every occurrence. The construction preserves differences in feature amplitudes but does not incorporate the nonzero Laplacian spectrum, which enters the Grassmann representation in the next subsection.

### 2.5 Grassmann chordal distance

To combine *k*-mer topology with Grassmannian geometry, take all four channels in (6). Normalise each block by a corpus-level root-mean-square scale *s*_*g,k*_ and an order weight *w*_*k*_ = 2^−(*K*−*k*)^, and stack the normalised blocks vertically over (*g, k*) into one matrix

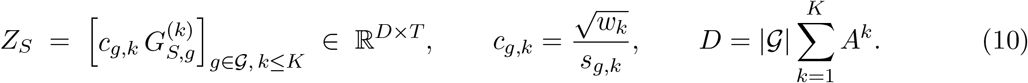

Compute a singular value decomposition of *Z*_*S*_ [31, Section 2.4] and let 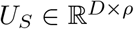collect the *ρ* leading left singular vectors. The representation of *S* is the subspace they span,

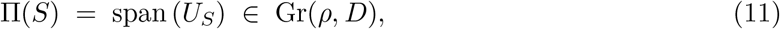

and the singular values are not retained as coordinates. The truncation is unique when the *ρ*th singular value is separated from the next one. This is the representation of Wang and Wei [23]. Once the normalisation constants are fixed, this map assigns each sequence a point in a common Grassmann manifold. The distance is then computed between subspaces rather than between feature vectors.

#### Definition 2

(Chordal distance). Let *θ*_1_, …, *θ*_*ρ*_ be the principal angles between Π(*S*) and Π(*R*) [31, Section 6.4.3]. Following Conway et al. [30], their chordal distance is

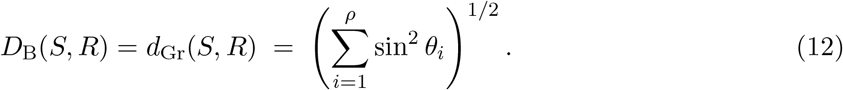

Since the singular values of 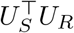are cos *θ*_*i*_, an equivalent expression for computation is

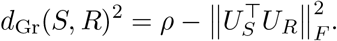

#### Proposition 3

(The chordal distance is a metric on Gr(*ρ, D*)). Let 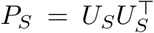denote the orthogonal projector onto Π(*S*), and let *θ*_1_, …, *θ*_*ρ*_ ∈ [0, *π/*2] be the principal angles between Π(*S*) and Π(*R*). Then

i. *d*_Gr_ is invariant under *U*_*S*_ ↦→ *U*_*S*_*O*_1_ and *U*_*R*_ ↦→ *U*_*R*_*O*_2_ for any *O*_1_, *O*_2_ ∈ *O*(*ρ*), so it is a well-defined function on Gr(*ρ, D*) *×* Gr(*ρ, D*);
ii. 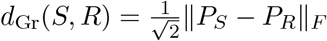, and *d*_Gr_ is a metric on Gr(*ρ, D*);
iii. 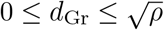, with *d*_Gr_ = 0 iff Π(*S*) = Π(*R*) and 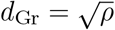iff the subspaces are orthogonal.

These standard projector and principal-angle expressions for the chordal metric are described by Conway et al. [30]; see also Edelman et al. [29].

The projector expression turns subspace comparison into a Euclidean distance between matrices; the angle expression describes the same distance through relative orientation. For rank-one subspaces, it reduces to 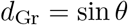. Neither expression depends on which orthonormal basis represents either subspace.

The two constructions compare different aspects of the sequence representation. The Betti-curve distance retains differences in feature amplitudes, whereas the Grassmann distance compares the orientations of leading subspaces and also incorporates non-harmonic spectral channels. For a well-defined truncation, uniform scaling *Z*_*S*_ ↦→ *cZ*_*S*_ with *c >* 0 leaves the Grassmann representation unchanged; reweighting individual channels can change it. Both constructions admit different alphabets, but their comparison also depends on the channels, radius grids and normalisation conventions. The chordal distance is a metric on the Grassmann manifold; on sequences, distinct inputs can share a representation and hence have zero distance. Thus the distinction is between comparing measured values in a feature space and comparing their leading subspaces; the present implementations also differ in the spectral measurements they include.

## 3 Data and methods

We combined HA sequences collected in the United States during August and September with historical clinical VE estimates to predict vaccine effectiveness for the upcoming influenza season. After sequence filtering, exclusion of nonseasonal viruses and verification of the published VE estimates, the final historical analysis included 1,226 HA sequences contributing to 26 observations (Supplementary Table S4) from 13 seasons between 2010–2011 and 2024–2025. Each observation paired the mean distance to the corresponding vaccine strain with one published VE estimate for a particular season and viral subtype or lineage. These 26 observations were used to fit the pooled regressions and to test predictions using earlier seasons to predict later ones. For the 2026–2027 forecasts, a separate set of 87 HA sequences collected in August–September 2026 passed the sequence filters: 44 A(H1N1)pdm09, 26 A(H3N2) and 17 B/Victoria. Their mean distances were used to generate forecasts from the historical regressions; no 2026–2027 VE outcome was used to fit or test the models. Figure 2 shows the procedure.

**Figure 2:**
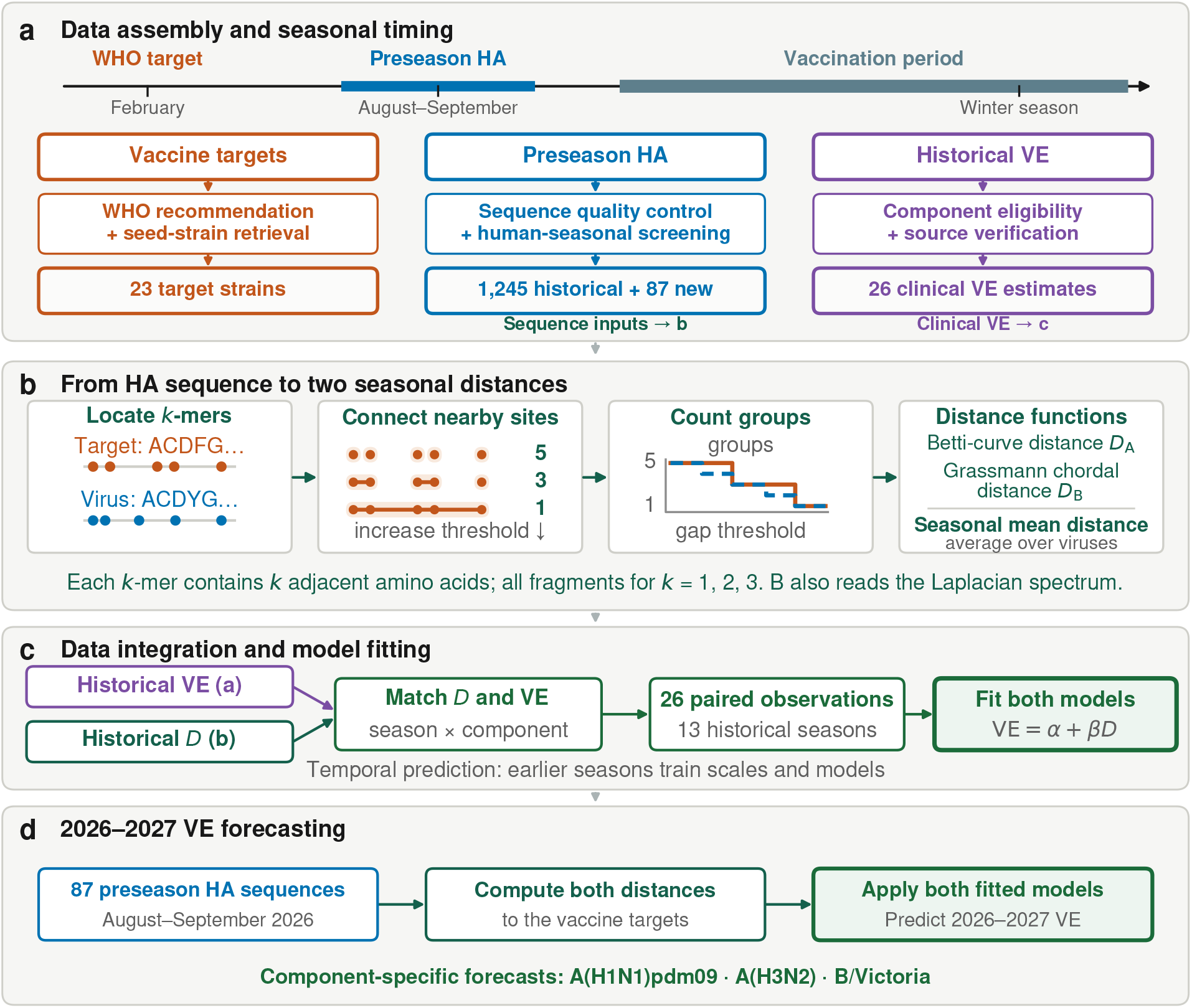
Data and prediction procedure. **(a)** Vaccine targets, preseason HA sequences and published component-specific VE estimates are assembled and screened. **(b)** *k*-mer position filtrations provide Betti-0 curves and Laplacian descriptors for the two distances, which are averaged by season and component. Sequence examples are schematic. **(c)** Distances are paired with 26 historical VE observations to fit two regressions; temporal validation fits scales and models on earlier seasons only. **(d)** The fitted models use distances from 87 August–September 2026 HA sequences to forecast 2026–2027 component-specific VE.

Each observation in the regression contains one mean distance and one VE estimate for a particular season and viral subtype or lineage. Individual sequences contribute to the mean distance but do not provide separate clinical outcomes. This distinction prevents a season with many submitted sequences from being counted as many independent VE observations.

### 3.1 HA sequences and vaccine strains

We obtained historical HA sequences from GISAID [21, 32, 33]. When downloading the data, we set the GISAID collection-date filter to August 1 through September 30 of each year, inclusive. The searches also required a United States location, a human host label and an available HA segment. This date filter refers to when the specimens were collected, rather than when the sequences were submitted or downloaded. The final historical analysis covered 13 seasons between 2010–2011 and 2024–2025. Supplementary Table S6 lists the GISAID dataset identifiers and DOIs for the searches.

We retained original-passage records with complete HA sequences of at least 1,650 nucleotides for influenza A and at least 1,700 nucleotides for influenza B, and excluded sequences containing ambiguous bases. We also checked for viruses that did not represent seasonal human influenza, because the host label entered by a submitting laboratory was not sufficient to establish this. Three rules, set before examining VE values, excluded records with a clade beginning with 1990, records assigned to clade 2010.1, or records with less than 95% gapped-alignment similarity to the corresponding vaccine strain. The first two rules identify swine-origin lineages; the sequence comparison also checks records with missing or unassigned clade labels. Each excluded accession and its reason were retained. These checks left 1,245 historical sequences, of which 1,226 belonged to the final 26 observations used in the VE analysis.

For comparison with the circulating viruses, we selected the vaccine strain recommended for the Northern Hemisphere in each season and for each viral subtype or lineage [34]. We obtained the corresponding HA sequences and checked strain names, sequence sources and passage histories before calculating distances. Supplementary Table S5 lists the vaccine strains and their sequence accessions.

We used the same search criteria and sequence filters to obtain the data for the 2026–2027 forecasts. These specimens were collected in the United States during August–September 2026. The searches were performed on 9 October 2026, after the collection period ended, and retained 87 HA sequences: 44 A(H1N1)pdm09, 26 A(H3N2) and 17 B/Victoria. We compared these sequences with the corresponding 2026–2027 vaccine strains and used their mean distances as inputs to the historical regression models. The GISAID dataset identifiers and DOIs for these searches are also listed in Supplementary Table S6.

### 3.2 Calculation of sequence distances

For the distance calculations, we used the HA protein of each circulating virus and vaccine strain, represented by its ordered sequence of amino acids. Both methods considered the 20 standard amino acids and recorded the positions of individual amino acids, adjacent pairs and adjacent triplets (*k* = 1, 2, 3). We calculated the two distances defined in Section 2 between each retained virus and the vaccine strain for the same season and subtype or lineage. For each method, we then averaged these virus–vaccine distances within each season and subtype or lineage. This average was the distance used to predict VE. Table 1 lists the settings for both methods.

**Table 1:** Settings for the two sequence distances. Both use the same HA sequences and vaccine strains. The listed scales apply to the historical pooled fits and frozen forecasts; prediction tests on later seasons estimate scales from earlier seasons only.

| | Topological distance $D_A$<br>($D_A, 20$) | Geometric distance $D_B$<br>($D_B, 23$) |
| --- | --- | --- |
| Alphabet | 20 amino acids | 20 amino acids |
| Orders $k$ | 1, 2, 3 | 1, 2, 3 |
| Channels $\mathcal{G}$ | $\beta_0$ | $\beta_0, \lambda_{\min}^+, \mu, \sigma$ |
| Radii | $\{1, 2, 4, \dots, N\}$, log<sub>2</sub>-spaced | $T = 50$, log-spaced on $[1, 600]$ |
| Order weights | $2^{-(K-k)}$ in (9) | $w_k = 2^{-(K-k)}$ in (10) |
| Scale | global $\omega_k$ (one per order) | global RMS $s_{g,k}$ (one per channel-order) |
| Rank $\rho$ | — | 10 |
| Comparison | Euclidean, pairwise | chordal on $\text{Gr}(10, D)$ |

The Betti-curve distance counted the connected groups of occurrences of each amino acid, pair or triplet at radii *r* ∈ *{*1, 2, 4, 8, …, *N}*, where *N* is the greater of the two sequence lengths. The Grassmann distance used these group counts together with the three Laplacian summaries defined in Section 2: the smallest positive eigenvalue, the mean of the positive eigenvalues and their standard deviation. It used 50 logarithmically spaced radii on [1, 600] and retained 10 leading singular vectors to represent each HA protein (*ρ* = 10). The upper radius exceeds the longest HA protein in the data, which contains 585 amino acids. Hozumi and Wei [20] sample radii linearly as *r*_*i*_ = 4 ^*k*−1^*i*, the factor 4 ^*k*−1^ keeping the number of filtration steps in proportion to the 4^*k*^ nucleotide *k*-mer types. With the 20-letter amino-acid alphabet the corresponding factor would advance the *k* = 3 radii in steps of several hundred residues, exceeding the length of an HA protein. We therefore sample logarithmically, which covers both short and long gaps along these proteins in a comparable number of steps. We considered individual amino acids, pairs and triplets because the 20^4^ = 160,000 possible combinations of four amino acids greatly outnumber the positions in an HA protein. The protein examples in Hozumi and Wei [20] also use *k* = 3.

We rescaled the measurements before combining the different word lengths and Laplacian summaries, so that measurements with larger numerical values would not dominate the distance calculations. For the historical pooled fits and the frozen forecasts, the Betti-curve scales were *ω*_1_ = 7.969, *ω*_2_ = 17.479 and *ω*_3_ = 24.826, calculated across the study comparisons. The Grassmann RMS scales *s*_*g,k*_ were fitted using all 529 distinct protein sequences, including those from 2026. Each resulting scale was shared across seasons and viral groups. Estimating a separate *ω*_*k*_ for each group would force its mean normalised distance to a fixed value and remove the differences needed for regression. The shared scales preserve those differences. Tests of prediction on later seasons used scales fitted only to earlier data, as described in the next subsection; the project-wide scales were not used for those tests.

We checked the Betti-curve calculation against the published formulas and internal consistency tests. For the Grassmann calculation, we used the reference implementation of Wang and Wei [23] for the features, normalisation, SVD and distance, using the 20 amino acids, word lengths of one to three and the 50 radii specified above. With the original nucleotide settings, the calculated feature blocks matched the reference results to machine precision. Further details of the settings and implementation checks are given in Supplementary Information, Section C.4.

### 3.3 VE prediction and evaluation

We matched the mean virus–vaccine distances for each season and subtype or lineage with the corresponding published clinical VE estimates [4–6, 35–44]. These estimates came primarily from the US Flu VE Network and used a test-negative design, which compares vaccination status between patients who test positive and those who test negative for influenza [2]. All of these estimates are adjusted estimates, computed as (1 − aOR) *×* 100 from multivariable logistic regression, where aOR is the adjusted odds ratio of testing positive for influenza among vaccinated compared with unvaccinated participants. The source articles adjust a priori for study site, participant age, presence of underlying health conditions or general health status, and month of illness onset, with some seasons additionally adjusting for race and ethnicity, so the covariate sets are similar but not identical across seasons. The quantity modelled here, and therefore the quantity forecast, is adjusted component-specific VE rather than any crude or unadjusted effectiveness. We checked each estimate against its source article and required it to match the subtype or lineage being analysed. The final 26 observations, including both mean distances, VE estimates, reported confidence intervals and source articles, are listed in Supporting Information, Section C.1 and Supplementary Table S4. The individual source checks, reasons for excluding two observations and checks of a figure-derived value are described in Supplementary Information, Section C.3.

For each distance, we fitted a separate pooled linear regression,

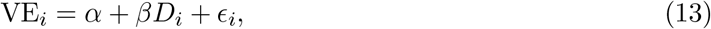

where *D*_*i*_ is the mean virus–vaccine distance for observation *i*, VE_*i*_ is the corresponding clinical VE in percentage points, *α* is the intercept, *β* is the slope and *ϵ*_*i*_ is the residual error. All subtypes and lineages entered the same regression with a common intercept and slope. We used ordinary least squares, giving each observation equal weight; sequence counts do not measure the precision of clinical VE.

We assessed the distance–VE association using regression standard errors and permutations of the VE labels. We report 95% OLS prediction intervals for future clinical VE under the regression model. Supplementary Information, Section C.5, describes the slope-permutation tests and prediction intervals.

We tested prediction by fitting the distance scales and regressions to earlier seasons and predicting the next test season. No sequence from the test season or a later season was used to estimate those scales. Both models were tested on the same observations and compared with a baseline that predicted the mean VE for the relevant subtype or lineage in earlier seasons. We measured absolute and squared prediction errors and the coverage and width of 95% prediction intervals. This retrospective evaluation follows season order rather than reconstructing historical publication and sequence-deposition dates. Supplementary Information, Section C.6, gives the test-season eligibility rules, distance-scale calculations and error measures.

The frozen 2026–2027 forecasts use the original project-wide scales and historical pooled regressions; they were not replaced by the later evaluation procedure. We will compare them with final US Flu VE Network estimates for the matching subtypes and lineage. As a separate check of the preseason measurements, we compared topological distances from August–September with those from the subsequent clinical enrollment periods for 18 matched observations across eight seasons. This comparison did not use VE values; its methods and data limitations are given in Supplementary Information, Section C.7.

## 4 Results

For the 2026–2027 influenza season in the United States, the pooled topological model predicted VE of 40.3% against A(H1N1)pdm09, 56.9% against A(H3N2) and 47.1% against B/Victoria. The pooled geometric model predicted 34.0%, 56.8% and 51.1%, respectively (Table 2). These are subtype- and lineage-specific point forecasts, not an overall VE estimate for the season, and their 95% prediction intervals are wide. Below, we first present the historical distance–VE fits used to obtain these forecasts, then compare August–September topological distances with distances measured during clinical enrollment, and report the new-season forecasts and their uncertainty. Prediction tests on later historical seasons are reported in Supplementary Information, Section B.2.

**Table 2:** Frozen 2026–2027 component forecasts from the pooled fit of each distance. The models use two distance representations of the same HA sequence collection, and their forecasts are reported separately. Intervals are 95% future-observation intervals. Both pooled forecasts are frozen and will be evaluated prospectively. No 2026–2027 VE outcome was used in fitting.

| Component | $n$ viruses | $k$-mer topological ($D_A$) | | | Grassmann geometric ($D_B$) | | |
| --- | --- | --- | --- | --- | --- | --- | --- |
|  |  | 2026–2027 |  |  | 2026–2027 |  |  |
| | | Mean $D$ | Predicted VE (%) | 95% PI | Mean $D$ | Predicted VE (%) | 95% PI |
| A(H1N1)pdm09 | 44 | 1.5804 | 40.3 | [13, 68] | 1.2706 | 34.0 | [8, 60] |
| A(H3N2) | 26 | 0.8506 | 56.9 | [28, 86] | 0.5497 | 56.8 | [29, 84] |
| B/Victoria | 17 | 1.2814 | 47.1 | [20, 75] | 0.7301 | 51.1 | [24, 78] |

### 4.1 Pooled relationships between sequence distance and VE

The historical analysis included 26 observations from 13 seasons, with both mean virus–vaccine distances paired with the same clinical VE estimate for each season and subtype or lineage (Supplementary Table S4). Larger distances were associated with lower VE for both methods (Figure 3). The fitted topological model was

**Figure 3:**
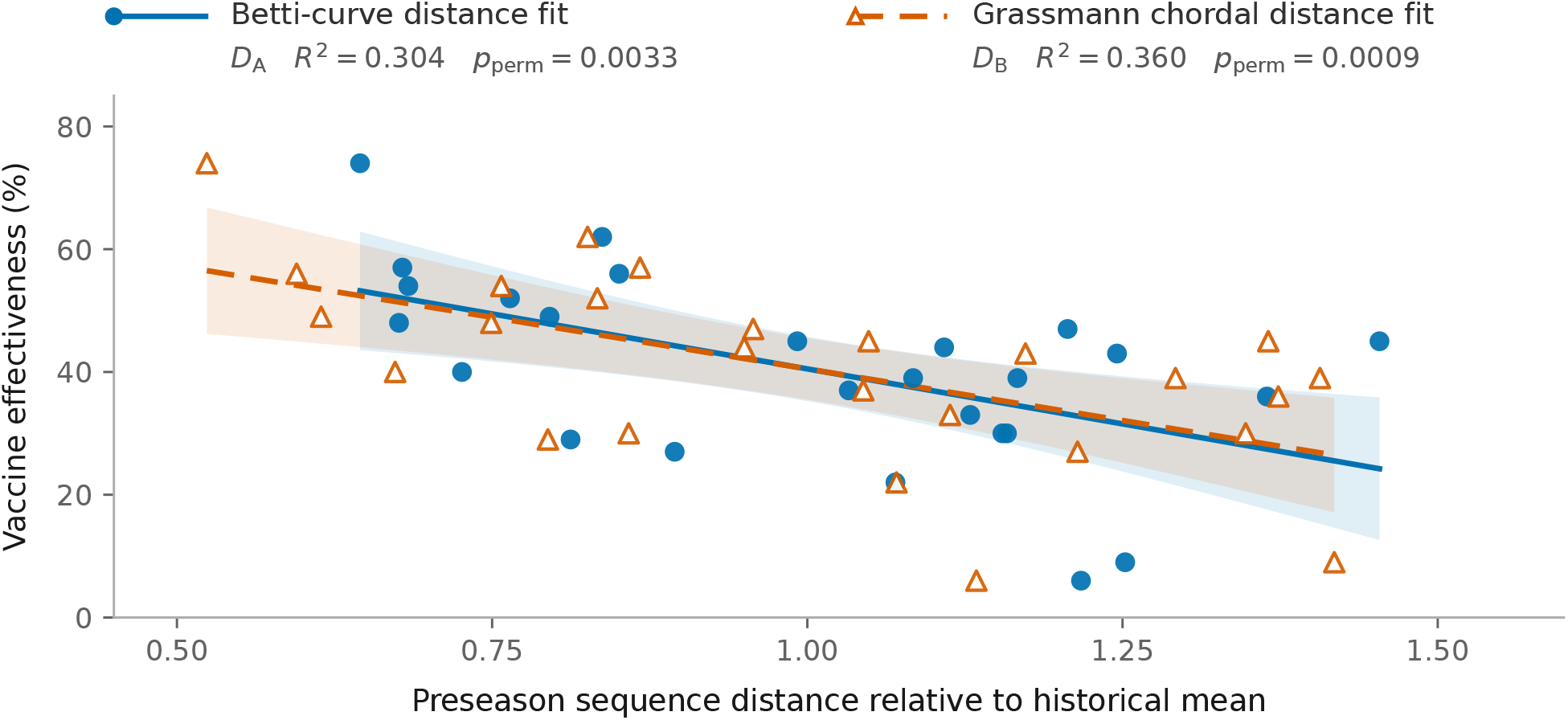
Pooled distance–VE fits. The 26 historical observations are shown for both methods: blue circles and a solid line for the Betti-curve distance *D*_*A*_; orange open triangles and a dashed line for the Grassmann chordal distance *D*_*B*_. Shading shows 95% confidence bands for mean VE. For display, each distance is divided by its historical mean (1 on the horizontal axis); this leaves fitted VE, *R*^2^ and *p* values unchanged. The slope-permutation *p* values test each association, not a difference between methods. These fits do not measure future prediction accuracy.

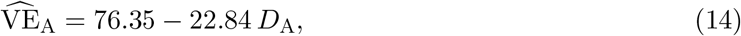

with *R*^2^ = 0.304 and a two-sided slope-permutation *p* = 0.0033. The fitted geometric model was

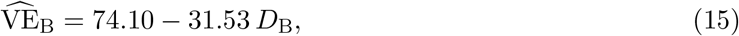

with *R*^2^ = 0.360 and a two-sided slope-permutation *p* = 0.0009. Thus, the models explained 30.4% and 36.0% of the variation in historical VE, respectively. These pooled fits describe associations across subtypes, lineages and seasons; they do not establish prediction accuracy for a future season.

The effects of excluding sequence records and VE observations on the pooled fit are reported in Supplementary Information, Section B.1. Component-adjusted analyses did not establish an association beyond subtype or lineage identity (Supplementary Information, Section A.1). Throughout, *component-adjusted* refers to the inclusion of subtype or lineage intercepts in the regression, which is distinct from the confounder adjustment already applied within each clinical VE estimate (Section 3.3). Separate fits within each subtype or lineage were also inconclusive (Section A.2; Figure S1).

### 4.2 August–September topological distances track enrollment-period distances

To use August–September sequences for an early forecast, we first examined whether their distances to the vaccine strain reflected those measured during the later clinical study. We compared topological distances for 18 matched observations from eight influenza seasons: 2014–2015, 2015– 2016, 2016–2017, 2017–2018, 2018–2019, 2019–2020, 2021–2022 and 2022–2023. Each observation paired the mean distance in August–September with that during enrollment for one season and subtype or lineage. The comparison included five A(H1N1)pdm09, six A(H3N2), four B/Victoria and three B/Yamagata observations. Seasons contributed one to four observations, depending on which viral groups had sequences available in both windows. Each pair used the same vaccine strain and shared distance scales. This allowed us to compare the earlier and later virus samples without changing the vaccine reference or the scale of the measurements.

The August–September and enrollment-period distances were strongly correlated (*r* = 0.942; Figure 4a). Observations with larger virus–vaccine distances before the season generally also had larger distances during clinical enrollment. However, this pooled correlation could partly reflect differences between subtypes or lineages that persist in both windows. We therefore subtracted each viral group’s mean separately in each window and compared the remaining deviations. Both panels show all 18 observations; subtracting the group means changes their coordinates without removing or combining observations. The correlation remained high (*r* = 0.844; Figure 4b): observations above or below their group’s mean in August–September tended to remain above or below it during enrollment. The season-cluster bootstrap 95% interval was 0.632–0.980, reflecting uncertainty across the eight available seasons.

**Figure 4:**
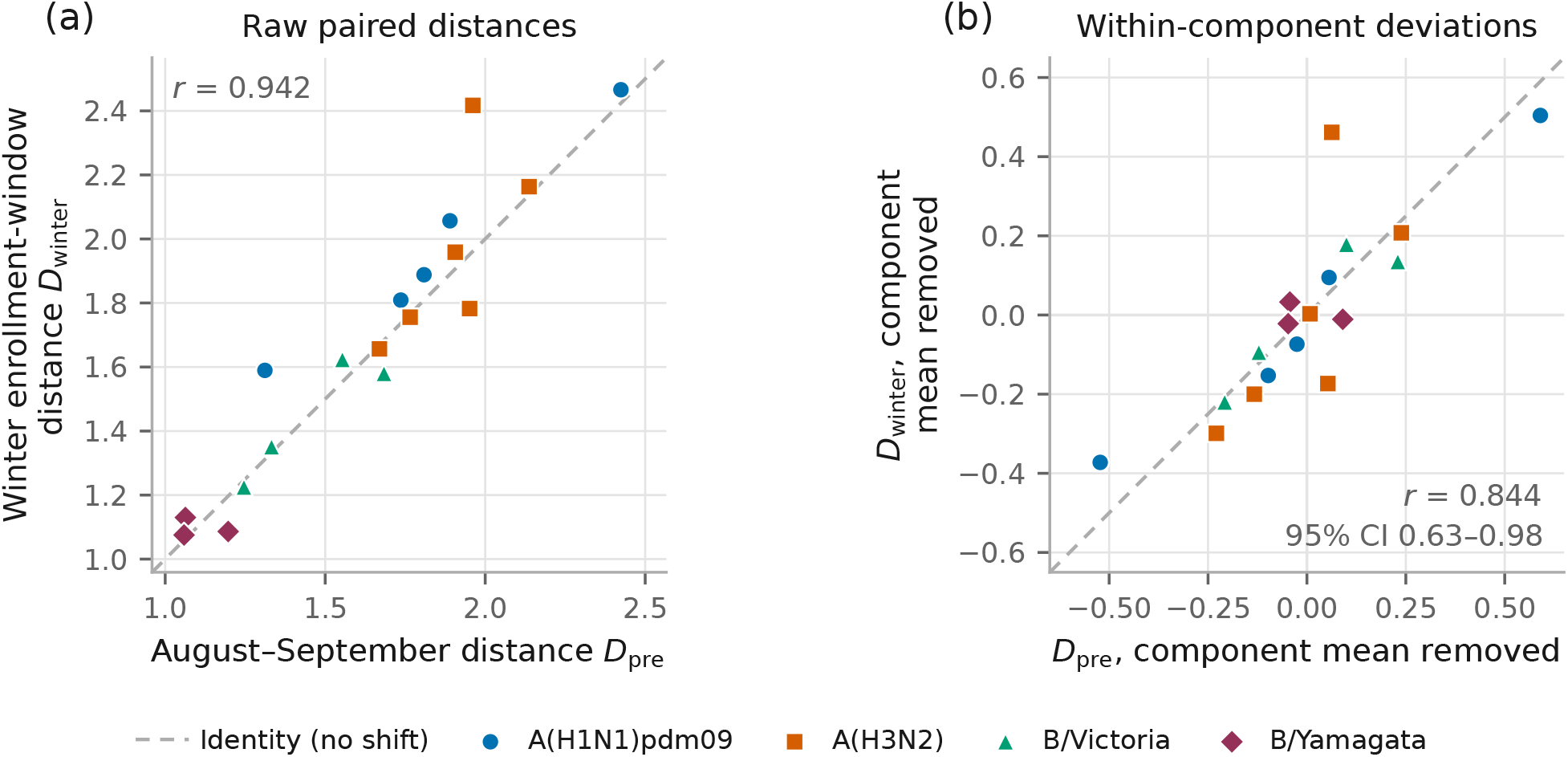
Preseason distance tracks winter enrollment-window distance. Both panels show the same 18 season–component observations from eight influenza seasons listed in Section 4.2: five A(H1N1)pdm09, six A(H3N2), four B/Victoria and three B/Yamagata. (a) Each point pairs the mean August–September distance with the mean enrollment-period distance for the same season and viral group. (b) The same 18 points after subtracting each viral group’s mean separately in each window, removing between-group differences in mean distance. No observations are removed or combined. The dashed line is identity in both panels. The 95% interval in (b) is a season-cluster bootstrap and remains a small-sample estimate. This figure establishes temporal concordance of the measurement, not its antigenic validity.

These results support using August–September topological distances as an early measure of the relative virus–vaccine differences observed during clinical enrollment. The relationship persisted after removing mean differences between viral groups, supporting the use of the earlier samples when enrollment-period sequences are not yet available. Correlation alone does not establish that the two windows give numerically interchangeable distances, however. The comparison was limited to seasons with data in both windows and was performed only for the topological distance. It does not directly validate the geometric distance or establish accurate clinical VE prediction. The matching procedure, shared scales and data limitations are described in Supplementary Information, Section C.7.

### 4.3 Pooled forecasts for the 2026–2027 season

The forecast data comprised 44 A(H1N1)pdm09, 26 A(H3N2) and 17 B/Victoria HA sequences from specimens collected during August–September 2026. The pooled topological model predicted VE of 40.3%, 56.9% and 47.1%, respectively; the pooled geometric model predicted 34.0%, 56.8% and 51.1% (Table 2). The two methods gave similar point forecasts, particularly for A(H3N2), and are reported separately without averaging.

The 95% prediction intervals were approximately 52–58 percentage points wide, reflecting substantial unexplained variation in the historical fits. Both sets of forecasts remain frozen for comparison with final 2026–2027 US Flu VE Network estimates for the matching subtypes and lineage.

## 5 Discussion

This study linked HA sequence distances to historical clinical VE and used the fitted relationships to forecast subtype- and lineage-specific VE for the 2026–2027 US influenza season. Both distances showed pooled associations with VE, and August–September topological distances tracked those measured during later clinical enrollment. Together, these findings support investigating preseason sequences as an early source of information for VE prediction. The limited improvement in historical prediction and wide forecast intervals nevertheless leave practical reliability to be established through prospective evaluation.

### Comparability of overall and component-specific VE estimates

The CDC past-seasons summary reports a single overall VE estimate against any influenza across all ages [45]. The forecasts here, however, concern individual subtypes and lineages. Both are adjusted clinical estimates in the sense of Section 3.3; the distinction is whether the outcome combines co-circulating viruses or refers to a particular viral group. This distinction matters when interpreting the forecasts and comparing them with subsequently reported clinical VE.

Table 3 compares the overall and component-specific estimates across the 13 seasons represented in this study. CDC overall VE exceeded the unweighted mean of the available component estimates by an average of 2.4 percentage points. Excluding 2024–2025 reduced this average difference to 0.8 percentage points. These small average differences conceal variation among viral groups. Overall VE exceeded the A(H3N2) estimate in 10 of 11 seasons and fell below the available B estimates in seven of eight seasons. This pattern is consistent with the lower VE commonly reported against A(H3N2) [1], but the available components do not represent every circulating group in every season. In particular, the four earliest seasons contributed only A(H3N2) observations. Their component means therefore equal the A(H3N2) estimates and cannot represent overall seasonal VE. The two largest negative differences, −7 in 2016–2017 and −17 in 2022–2023, coincide with A(H3N2) predominance and reduced vaccine match from egg-adapted vaccine strains in the former [40, 46], and with an early season following minimal influenza circulation during the COVID-19 pandemic in the latter [44, 47].

**Table 3:**
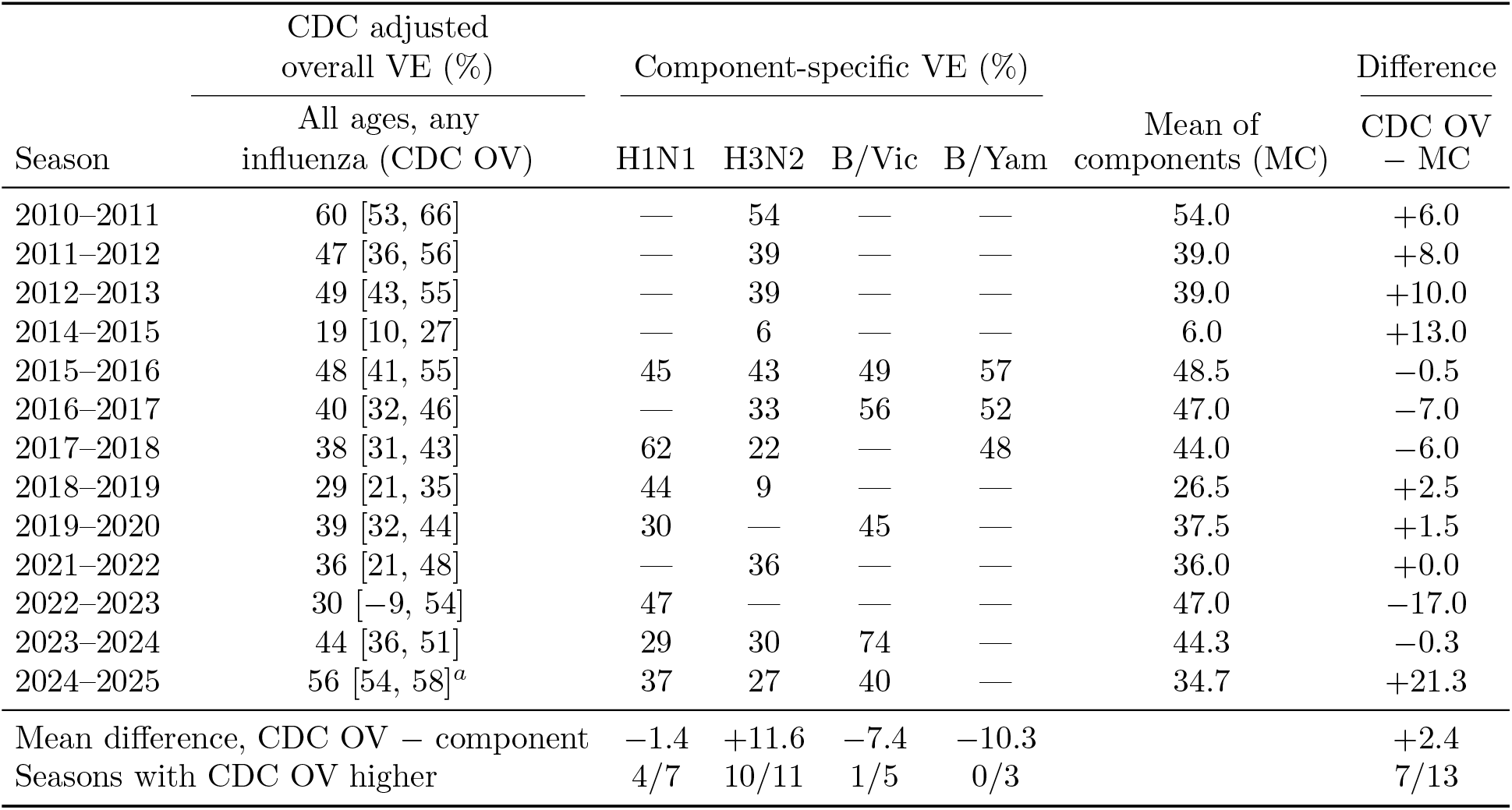
The CDC overall VE estimate for each season compared with the component-specific estimates used in this study. CDC values are adjusted overall VE (CDC OV) against any influenza across all ages, with 95% confidence intervals, as listed on the CDC past-seasons page [45]. Component values are the clinical estimates in Table S4; a dash indicates that no observation for that group was included for the season. The mean of the components (MC) is the unweighted average of the available group estimates and should not be interpreted as an estimate of overall seasonal VE. Differences are expressed in percentage points. ^*a*^The 2024–2025 CDC entry comes from the VISION network, whereas the component estimates come from the US Flu VE Network [48, 49].

| Season | CDC adjusted<br>overall VE (%) | Component-specific VE (%) |  |  |  | Mean of<br>components (MC) | Difference<br>CDC OV<br>– MC |
| --- | --- | --- | --- | --- | --- | --- | --- |
|  | All ages, any<br>influenza (CDC OV) | H1N1 | H3N2 | B/Vic | B/Yam |  |  |
| 2010–2011 | 60 [53, 66] | — | 54 | — | — | 54.0 | +6.0 |
| 2011–2012 | 47 [36, 56] | — | 39 | — | — | 39.0 | +8.0 |
| 2012–2013 | 49 [43, 55] | — | 39 | — | — | 39.0 | +10.0 |
| 2014–2015 | 19 [10, 27] | — | 6 | — | — | 6.0 | +13.0 |
| 2015–2016 | 48 [41, 55] | 45 | 43 | 49 | 57 | 48.5 | –0.5 |
| 2016–2017 | 40 [32, 46] | — | 33 | 56 | 52 | 47.0 | –7.0 |
| 2017–2018 | 38 [31, 43] | 62 | 22 | — | 48 | 44.0 | –6.0 |
| 2018–2019 | 29 [21, 35] | 44 | 9 | — | — | 26.5 | +2.5 |
| 2019–2020 | 39 [32, 44] | 30 | — | 45 | — | 37.5 | +1.5 |
| 2021–2022 | 36 [21, 48] | — | 36 | — | — | 36.0 | +0.0 |
| 2022–2023 | 30 [–9, 54] | 47 | — | — | — | 47.0 | –17.0 |
| 2023–2024 | 44 [36, 51] | 29 | 30 | 74 | — | 44.3 | –0.3 |
| 2024–2025 | 56 [54, 58]<sup>a</sup> | 37 | 27 | 40 | — | 34.7 | +21.3 |
| Mean difference, CDC OV – component |  | –1.4 | +11.6 | –7.4 | –10.3 |  | +2.4 |
| Seasons with CDC OV higher |  | 4/7 | 10/11 | 1/5 | 0/3 |  | 7/13 |

Differences in surveillance network also affect the comparison. The 2024–2025 overall entry on the CDC summary page comes from the VISION network, whereas the component estimates used here come from the US Flu VE Network [48, 49]. The overall estimates from the two networks differed by 14 percentage points in that season. Such differences reinforce the need to match the clinical setting, study population and viral group when evaluating a forecast.

We therefore specify the intended clinical comparison in advance. The frozen forecasts are to be evaluated against final US Flu VE Network adjusted estimates for medically attended outpatient illness with laboratory-confirmed influenza, separately for A(H1N1)pdm09, A(H3N2) and B/Victoria. Evaluation should consider both the point predictions and their prediction intervals. Comparing these forecasts directly with a single overall VE estimate would not assess the outcomes the models were fitted to predict.

### Interpreting the pooled associations

The fitted relationships combine differences across subtypes, lineages and seasons. They do not establish that changes in distance within each viral group correspond to changes in VE. After accounting for differences in mean VE among groups, the shared distance slope was not statistically significant (Supplementary Information, Section A.1; Supplementary Table S1). Separate fits for individual subtypes and lineages were also inconclusive (Section A.2; Figure S1). The pooled association may therefore reflect both between-group differences and seasonal variation within groups, whose contributions remain unresolved. The small number of observations within each group also limits the interpretation of nonsignificant slopes.

The window comparison addresses a different question: whether earlier samples reflect relative virus–vaccine differences during clinical enrollment. Its positive result supports the timing of the measurement in the available seasons, but does not independently validate the relationship with clinical VE. Agreement varied across groups and seasons and was assessed only for the topological distance. Detailed influence checks and group-specific results are provided in Supplementary Information, Section C.7.

### Historical prediction accuracy and uncertainty

Fitting distance scales and regression coefficients using earlier seasons provides a more direct assessment of forecasting performance than the pooled fit alone. Across 19 observations from nine test seasons, mean absolute errors were 12.01 percentage points for the topological model and 11.34 for the geometric model. The reference that predicted each component’s earlier mean VE had an error of 11.96 percentage points. The geometric model’s improvement over this reference was therefore 0.62 percentage points. Mean absolute error and root mean squared error also ranked the two distance models differently. These results indicate limited improvement over the reference and do not demonstrate a consistent advantage for either distance representation.

Prediction intervals further indicate substantial uncertainty around individual VE forecasts. Their coverage must be considered together with their width, because wide intervals can contain observed VE while offering limited precision. Prediction errors, interval coverage and interval widths are reported in Supplementary Information, Section B.2 and Supplementary Table S2. The evaluation followed season order without fully reconstructing the information available at historical forecast dates. Observations from the same season also share a fitted model and should not be treated as independent seasonal replications.

The two methods differ in their sequence summaries, radius settings and normalisation, as described in Section C.4. Their performance differences therefore cannot be attributed to Grassmann geometry alone. Prospective evaluation against the specified clinical outcomes will help determine whether either representation provides useful predictive information beyond historical VE.

## 6 Conclusion

Topological and geometric distances between August–September HA sequences and vaccine strains were associated with historical clinical VE across 26 season–component observations. The topological distances also tracked those measured during later clinical enrollment in the available paired seasons, supporting their use as an early measure of relative virus–vaccine differences. Using the historical pooled relationships, we generated frozen 2026–2027 US VE forecasts of 40.3%, 56.9% and 47.1% for A(H1N1)pdm09, A(H3N2) and B/Victoria, respectively, with the topological model, and 34.0%, 56.8% and 51.1% with the geometric model. These are component-specific point forecasts with wide prediction intervals. Historical prediction gains were limited, and information beyond subtype or lineage identity was not well established. Comparison with the corresponding final clinical VE estimates, together with evaluation on additional independent data, is needed to determine whether these sequence-based forecasts can reliably support early-season planning.

## Data and code availability

Analysis code, record-level exclusion reasons with accessions, aggregated model inputs, and the saved forecasts and figure data are available at https://github.com/XiangXiangJY/Antigenic. GISAID sequences are subject to GISAID data-access terms and are identified by EPI SET accessions and DOIs (Supplementary Table S6) rather than redistributed. Every preseason search contributing to the cohort or the forecast carries an EPI SET identifier (Supplementary Table S6). The 2022–2023 A(H1N1)pdm09 identifier was generated on 14 October 2026 by re-running the original search specification, which returned exactly the same 99 records as the original download.

## Acknowledgements

We acknowledge the authors, originating laboratories and submitting laboratories that contributed the specimens, sequences and metadata shared through GISAID (Supplementary Table S6). This work was supported in part by NIH grant R01AI164266, the University of Georgia (UGA), and the Georgia Research Alliance.

## Supplementary Information

### A Component-adjusted and within-group associations

The pooled fits combine observations from different subtypes and lineages. Here, we examine whether distance remains associated with VE after allowing separate group intercepts, and whether a relationship is evident within each group separately. Neither analysis established a significant distance–VE association. These results limit the interpretation of the pooled associations, while the small group sizes leave the within-group relationships uncertain.

#### A.1 Different subtype and lineage intercepts with a shared distance slope

To assess whether distance provided information beyond subtype or lineage, we fitted

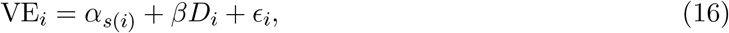

where *s*(*i*) identifies the subtype or lineage of observation *i*. Each of A(H1N1)pdm09, A(H3N2), B/Victoria and B/Yamagata has a separate intercept *α*_*s*(*i*)_, while *β* is a single distance slope shared by all four groups. Thus, the model allows different VE levels between groups and estimates the distance–VE relationship after accounting for these mean differences. The topological slope was −13.40 (SE 9.32; 95% CI, −32.8 to 6.0; *p* = 0.1651; model *R*^2^ = 0.4420; Table S1). The geometric slope was −17.43 (SE 12.11; 95% CI, −42.6 to 7.8; *p* = 0.1649; model *R*^2^ = 0.4421). The two adjusted fits happen to agree closely on both quantities. Neither adjusted slope was significant.

Adjustment reduced the magnitude of the topological slope from −22.84 to −13.40 and the geometric slope from −31.53 to −17.43. This attenuation is consistent with part of the pooled association reflecting differences between viral groups. Both adjusted confidence intervals included zero, so these data did not establish that distance contributed information about VE beyond subtype or lineage identity. However, the wide intervals also allow substantial negative slopes; they do not demonstrate that the within-group relationship is absent.

The adjusted models had higher *R*^2^ than the pooled models, but the extra group intercepts also allow a closer fit to the same observations. This increase describes overall fit and does not establish an additional contribution from distance or improved prediction on later seasons. The common-slope assumption also means that this analysis does not test whether the distance–VE relationship differs between groups; separate exploratory fits are described in Section A.2.

Excluding the figure-derived 2017–2018 B/Yamagata VE changed the adjusted topological slope from −13.40 (*t* = −1.44) to −13.60 (*t* = −1.43). The conclusion was therefore unchanged when this observation was omitted.

#### A.2 Separate distance–VE fits for each subtype and lineage

We examined whether the pooled distance–VE relationship was also evident across seasons within individual subtypes or lineages. For this exploratory analysis, we fitted a separate intercept and slope for each viral group, rather than imposing the common slope used in Section A.1. The historical dataset contained seven A(H1N1)pdm09, eleven A(H3N2), five B/Victoria and three B/Yamagata observations.

**Table S1:** Topological pooled and component-adjusted regression estimates on 26 observations. Standard errors are ordinary OLS.

| Term | Estimate | SE | 95% CI | OLS $p$ | $R^2$ |
| --- | --- | --- | --- | --- | --- |
| **Primary: pooled** |  |  |  |  |  |
| Intercept | 76.35 | 11.35 |  |  |  |
| Distance $D$ | -22.84 | 7.05 | [-37.4, -8.3] | 0.003 | 0.304 |
| **Sensitivity: subtype fixed effects** |  |  |  |  |  |
| Distance $D$ | -13.40 | 9.32 | [-32.8, 6.0] | 0.165 | 0.442 |
| Intercept: A(H1N1)pdm09 | 64.86 | 16.57 |  |  |  |
| Intercept: A(H3N2) | 54.18 | 16.73 |  |  |  |
| Intercept: B/Victoria | 69.67 | 12.98 |  |  |  |
| Intercept: B/Yamagata | 67.19 | 12.57 |  |  |  |

Within-group analyses did not establish a distance–VE association for any subtype or lineage with either distance. Figure S1a–d shows the topological results. All four topological slope estimates were negative, but their 95% confidence intervals included zero. The small group sizes and wide intervals leave the within-group relationships unresolved. These fits assess whether the pooled association also appears within individual groups and were not used to generate the 2026–2027 forecasts. In particular, the nearly flat H1N1 estimate and the narrow range of B/Yamagata distances provide little evidence of a common pattern within every group. Negative point estimates alone do not establish such a pattern.

**Figure S1:**
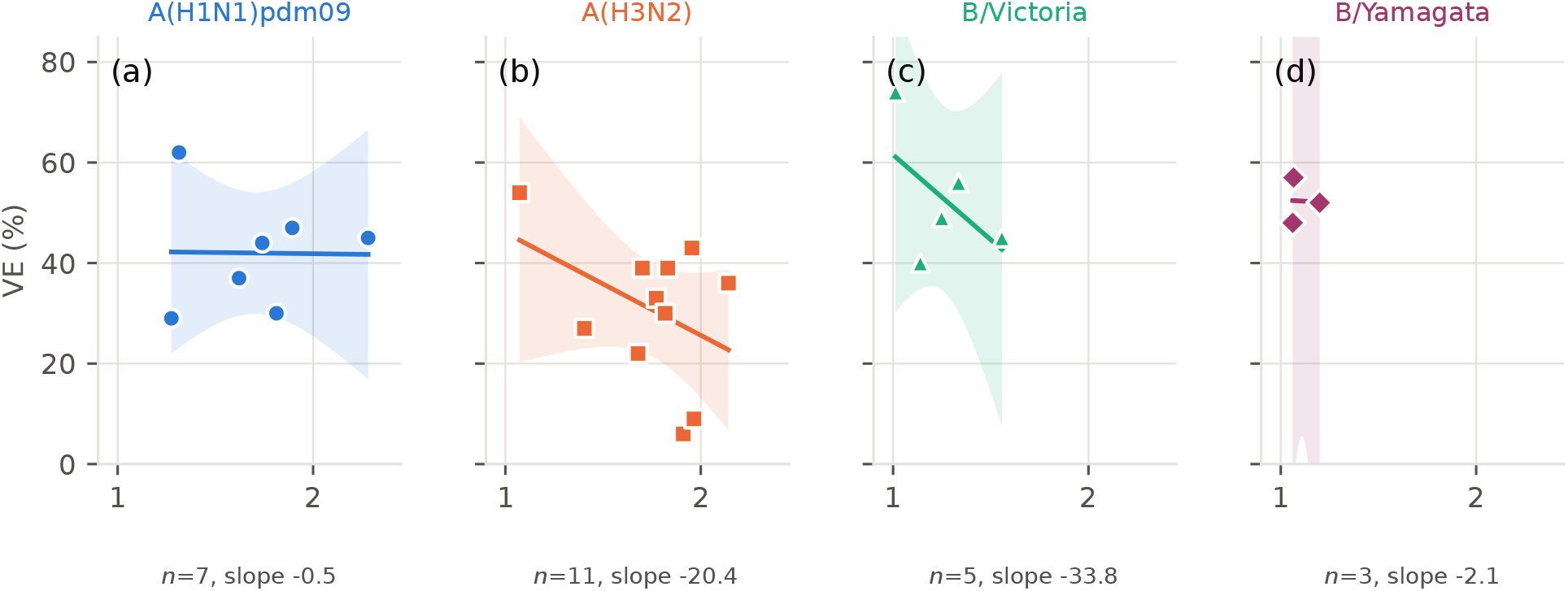
Separate distance–VE fits for each subtype and lineage. (a–d) Historical observations for A(H1N1)pdm09 (*n* = 7), A(H3N2) (*n* = 11), B/Victoria (*n* = 5) and B/Yamagata (*n* = 3), respectively. In every panel, the horizontal axis is the mean August–September topological distance *D*_*A*_ to the vaccine strain, and the vertical axis is clinical VE (%). Each group has a separately estimated intercept and slope. Solid lines and shading show ordinary least-squares fits and 95% confidence bands for mean VE in all four panels. B/Yamagata has only three observations spanning a distance range of 0.14, resulting in a very wide confidence band that extends beyond the displayed VE range. All four slope estimates are negative, but their 95% confidence intervals include zero. These exploratory fits were not used for forecasting.

### B Pooled model fits and predictions for later seasons

This section examines how excluding sequence records and VE observations affects the pooled fits, then evaluates predictions for later seasons using only earlier seasons for training. The fitted relationships changed after data exclusions. In historical prediction tests, the geometric pooled model modestly reduced absolute error relative to the historical-mean reference, but prediction intervals remained wide.

#### B.1 How sequence and VE exclusions changed the pooled fit

Sequence quality checks left 1,285 historical and 87 forecast-season records. The seasonal-human-virus checks excluded 40 historical records: 15 by the clade-prefix rule, 19 by clade 2010.1 and six by the similarity rule when applied in that order. Independently, the similarity rule flagged all 40. Of 1,245 retained historical sequences, 1,226 contributed to the final 26 observations (7 H1N1, 11 H3N2, five B/Victoria and three B/Yamagata). Clinical source checks reduced 28 assembled observations to 26; the two exclusions were fixed before refitting (Section C.3). They changed the pooled topological slope from −19.26 (*R*^2^ = 0.190) to −22.84 (*R*^2^ = 0.304).

With the same 26 clinical outcomes, using the sequence sets before the seasonal-human-virus exclusions gave slope −9.02 (*R*^2^ = 0.131; slope-permutation *p* = 0.075), compared with −22.84 (*R* = 0.304; *p* = 0.0033) after exclusions. Six group means changed; the largest changes were in 2012–2013 H3N2 (11 of 12 records removed) and 2010–2011 H3N2 (two of five removed). These changes show that the pooled fit depends materially on which sequence records and clinical estimates enter the analysis. Its interpretation therefore rests on the documented eligibility rules and source checks. Table S4 provides the final paired data.

#### B.2 Historical prediction performance of the pooled models

We assessed whether the pooled models could predict VE for a later season using distances and VE observations from earlier seasons. At each test season, distance scales and regression coefficients were re-estimated from the earlier seasons only. Both models were evaluated on the same 19 season–component observations across nine test seasons. Each pooled model was refitted nine times, once per test season. The fit for a season produced predictions for all eligible viral groups in that season. Each prediction corresponds to one clinical VE estimate for a particular season and subtype or lineage. The eligible seasons and inclusion rules are specified in Section C.6.

Table S2 compares prediction errors with a reference that uses the earlier mean VE for the relevant subtype or lineage. This reference asks whether sequence distances add predictive value beyond previous VE levels. The geometric pooled MAE was 11.34 percentage points and the topological pooled MAE was 12.01, against 11.96 for the reference. Both models had lower RMSE than the reference, the topological model the lower of the two.

These summaries rank the approaches differently. The topological model had the slightly higher mean absolute error (12.01 against 11.96) yet was closer to the observed VE in 11 of the 19 tests, against 8 for the reference, a split that an exact sign test does not distinguish from chance (*p* = 0.65); its absolute errors also had the smallest spread of the three (standard deviation 7.97 percentage points, against 9.56 for the geometric model and 9.69 for the reference). The higher mean therefore reflects a few large errors rather than a systematic difference. None of the paired differences in absolute error was distinguishable from zero: topological minus reference +0.05 percentage points (paired bootstrap 95% interval −2.97 to +3.12), geometric minus reference −0.62 (−2.61 to +1.52), and geometric minus topological −0.67 (−3.58 to +2.53). With 19 tests, a split of at least 15 to 4 would be needed to reach the conventional significance threshold. These seasons therefore cannot establish an ordering among the two distances and the reference.

The 95% prediction intervals contained the observed VE in 19 of 19 tests for the topological model and 18 of 19 for the geometric model. Their mean widths were 61.0 and 63.0 percentage points, respectively. High coverage therefore accompanied wide uncertainty about individual VE outcomes. Multiple observations came from the same season, and the evidence spans only nine test seasons. These results describe retrospective prediction performance and do not establish reliable prospective forecasts.

**Table S2:** Prediction accuracy and interval coverage for 19 historical VE observations across nine test seasons. Every row uses the same 19 season–component observations. Each pooled model was refitted for each test season using earlier seasons only. MAE is mean absolute error; RMSE is root mean squared error. Both errors and interval widths are in percentage points (pp). “Observed VE inside PI” counts how many actual VE estimates fall within their predicted 95% intervals. Coverage is this count divided by 19, expressed as a percentage. Thus, the geometric model made 19 predictions, and 18 observed VE estimates fell inside its intervals.

| Model or reference | Total predictions | Prediction error |  | 95% prediction intervals (PI) |  |  |
| --- | --- | --- | --- | --- | --- | --- |
|  |  | MAE (pp) | RMSE (pp) | Observed VE inside PI (count) | Coverage (%) | Mean width (pp) |
| Historical-mean reference | 19 | 11.96 | 15.23 | 19 | 100.0 | 77.9 |
| Topological pooled | 19 | 12.01 | 14.30 | 19 | 100.0 | 61.0 |
| Geometric pooled | 19 | 11.34 | 14.67 | 18 | 94.7 | 63.0 |

Table S3 lists the 19 individual predictions behind these summaries. Absolute errors ranged from 2.5 to 30.1 percentage points for the topological model and from 0.9 to 38.3 percentage points for the geometric model, and both models predicted a higher VE than was observed in 11 of the 19 tests. The geometric prediction was closer to the observed VE in 13 of the 19 tests, but its single largest error was larger than any topological error; the two error measures therefore rank the models differently. Each model was closer to the observed VE than the historical-mean reference in about half of the tests, 11 of 19 for the topological model and 10 of 19 for the geometric model, so the small differences in average error do not correspond to a consistent advantage on individual season–component outcomes.

The largest error for all three approaches occurred in the 2014–2015 A(H3N2) test, which had the lowest observed VE in the data (6%) and was also the earliest test season, fitted on only three earlier observations. This test is the one case in which an observed VE fell outside a prediction interval, the lower limit of the geometric interval being 12.3%. Individual interval widths ranged from 53.9 to 79.2 percentage points for the topological model and from 55.4 to 91.4 for the geometric model. Predictions for single season–component outcomes are therefore compatible with a wide range of VE values, and the historical-mean reference intervals are wider still, extending below zero and above 100% in several early tests.

### C Sequence records, VE sources and calculation procedures

This section documents the data and calculations behind the reported results. It first lists the 26 distance–VE observations used to fit the models, followed by vaccine strain records, GISAID sequence sets and checks of the published VE estimates. The remaining subsections describe distance calculations, statistical tests, prediction evaluation and the matching of sampling windows. They also identify limitations in record availability, clinical estimates and the coverage of the window comparison.

**Table S3:** Individual predictions for the 19 historical test observations summarised in Table S2. Each row is one clinical VE estimate for one season and subtype or lineage. “Training observations” is the number of earlier season–component observations used to re-estimate the distance scales and regression coefficients for that test season; all rows from the same test season share the same fit. Predictions and intervals are in VE percentage points, and the intervals are 95% future-observation intervals from the fit for that season. They are not constrained to the 0–100% range, so some reference limits fall outside it. ^*†*^ marks the one prediction whose interval does not contain the observed VE. The final rows repeat the summary measures of Table S2, recomputed from the rows shown here.

| Test season | Subtype or lineage | Training observations | Observed VE (%) | Predicted VE (95% prediction interval) |  |  |
| --- | --- | --- | --- | --- | --- | --- |
|  |  |  |  | Topological pooled | Geometric pooled | Historical-mean reference |
| 2014–2015 | A(H3N2) | 3 | 6 | 36.1 (3.3, 68.9) | 44.3 (12.3, 76.2)<sup>†</sup> | 44.0 (1.0, 87.0) |
| 2016–2017 | A(H3N2) | 8 | 33 | 38.9 (1.2, 76.6) | 39.7 (0.2, 79.1) | 36.2 (−18.4, 90.8) |
| 2016–2017 | B/Victoria | 8 | 56 | 47.0 (8.5, 85.5) | 53.5 (7.8, 99.2) | 49.0 (−21.5, 119.5) |
| 2016–2017 | B/Yamagata | 8 | 52 | 49.5 (9.9, 89.1) | 46.5 (6.0, 86.9) | 57.0 (−13.5, 127.5) |
| 2017–2018 | A(H1N1)pdm09 | 11 | 62 | 48.6 (19.0, 78.2) | 46.7 (16.5, 77.0) | 45.0 (−1.2, 91.2) |
| 2017–2018 | A(H3N2) | 11 | 22 | 40.9 (11.8, 70.0) | 40.7 (10.6, 70.8) | 35.7 (0.4, 71.0) |
| 2017–2018 | B/Yamagata | 11 | 48 | 53.9 (22.8, 85.0) | 49.3 (18.5, 80.1) | 54.5 (14.5, 94.5) |
| 2018–2019 | A(H1N1)pdm09 | 14 | 44 | 38.5 (9.7, 67.3) | 43.0 (13.7, 72.3) | 53.5 (18.2, 88.8) |
| 2018–2019 | A(H3N2) | 14 | 9 | 33.4 (3.7, 63.2) | 27.7 (−4.9, 60.4) | 33.7 (2.9, 64.5) |
| 2019–2020 | A(H1N1)pdm09 | 16 | 30 | 34.7 (4.9, 64.6) | 47.7 (19.1, 76.3) | 50.3 (15.8, 84.9) |
| 2019–2020 | B/Victoria | 16 | 45 | 41.5 (12.0, 71.0) | 26.1 (−3.8, 56.0) | 52.5 (15.8, 89.2) |
| 2021–2022 | A(H3N2) | 18 | 36 | 25.7 (−3.4, 54.8) | 28.6 (−2.1, 59.3) | 30.6 (−0.4, 61.7) |
| 2022–2023 | A(H1N1)pdm09 | 19 | 47 | 33.5 (6.3, 60.7) | 43.2 (14.8, 71.7) | 45.2 (13.7, 76.8) |
| 2023–2024 | A(H1N1)pdm09 | 20 | 29 | 49.1 (21.7, 76.4) | 47.5 (19.6, 75.4) | 45.6 (15.8, 75.4) |
| 2023–2024 | A(H3N2) | 20 | 30 | 36.3 (9.4, 63.2) | 30.9 (2.4, 59.3) | 31.2 (2.5, 59.9) |
| 2023–2024 | B/Victoria | 20 | 74 | 55.2 (26.7, 83.7) | 56.5 (26.6, 86.4) | 50.0 (18.6, 81.4) |
| 2024–2025 | A(H1N1)pdm09 | 23 | 37 | 40.5 (12.9, 68.2) | 40.2 (12.5, 67.9) | 42.8 (13.0, 72.6) |
| 2024–2025 | A(H3N2) | 23 | 27 | 46.0 (18.2, 73.9) | 34.0 (6.0, 62.0) | 31.1 (2.2, 60.0) |
| 2024–2025 | B/Victoria | 23 | 40 | 52.8 (24.2, 81.3) | 52.6 (24.1, 81.1) | 56.0 (25.1, 86.9) |
| Mean absolute error (pp) |  |  |  | 12.01 | 11.34 | 11.96 |
| Root mean squared error (pp) |  |  |  | 14.30 | 14.67 | 15.23 |
| Observed VE inside interval |  |  |  | 19/19 | 18/19 | 19/19 |
| Mean interval width (pp) |  |  |  | 61.0 | 63.0 | 77.9 |

#### C.1 Historical distances and VE estimates used to fit the models

Table S4 lists the 26 observations used to fit the historical VE models. Each row represents one season and subtype or lineage and pairs its mean virus–vaccine distance with a published clinical VE estimate. The two distance columns were calculated from the same sequence records; the sequence count identifies how many viruses contributed to each mean. Clinical confidence intervals describe uncertainty in the published VE estimates but were not used as regression weights. The source checks and exclusions are explained in Section C.3; their effect on the pooled fit is reported in Section B.1.

#### C.2 Vaccine strain records and GISAID sequence sets

Tables S5 and S6 identify the vaccine targets and registered preseason sequence sets. The 23 distinct seed strains supply 43 season–component assignments. Five assignments use records deposited after the corresponding season: B/Victoria in 2010–2011, 2011–2012, 2014–2015 and 2015–2016, and H3N2 in 2012–2013. Although strain names match the recommendations, these later records limit strict claims about historical data availability. Target mapping was checked manually.

**Table S4:** Historical distance–VE data used to fit the models (26 observations). Mean *D*_*A*_ and *D*_*B*_ are the original-scale Betti-curve and Grassmann distances, rounded to four decimal places, calculated from the same *n* viruses against the corresponding vaccine strain. VE estimates and references identify the clinical outcomes; 95% CIs are recorded where available and do not enter the regression. Source-specific qualifications are given in Section C.3.

| Season | Component | $n$ viruses | Mean $D_A$ | Mean $D_B$ | VE (%) | 95% CI | VE source |
| --- | --- | --- | --- | --- | --- | --- | --- |
| 2010–2011 | A(H3N2) | 3 | 1.0728 | 0.8070 | 54 | [42, 64] | [35] |
| 2011–2012 | A(H3N2) | 4 | 1.8310 | 1.4990 | 39 | [23, 52] | [36] |
| 2012–2013 | A(H3N2) | 1 | 1.7012 | 1.3769 | 39 | [29, 47] | [37] |
| 2014–2015 | A(H3N2) | 10 | 1.9102 | 1.2086 | 6 | [-5, 17] | [38] |
| 2015–2016 | A(H1N1)pdm09 | 8 | 2.2818 | 1.1175 | 45 | — | [39] |
| 2015–2016 | A(H3N2) | 209 | 1.9550 | 1.2501 | 43 | — | [39] |
| 2015–2016 | B/Victoria | 5 | 1.2487 | 0.6547 | 49 | — | [39] |
| 2015–2016 | B/Yamagata | 18 | 1.0654 | 0.9243 | 57 | — | [39] |
| 2016–2017 | A(H3N2) | 99 | 1.7723 | 1.1863 | 33 | [23, 41] | [40] |
| 2016–2017 | B/Victoria | 12 | 1.3353 | 0.6340 | 56 | [23, 75] | [40] |
| 2016–2017 | B/Yamagata | 12 | 1.1994 | 0.8882 | 52 | [42, 61] | [40] |
| 2017–2018 | A(H1N1)pdm09 | 33 | 1.3141 | 0.8800 | 62 | [50, 71] | [4] |
| 2017–2018 | A(H3N2) | 166 | 1.6792 | 1.1411 | 22 | [12, 31] | [4] |
| 2017–2018 | B/Yamagata | 38 | 1.0612 | 0.7985 | 48 | — | [4] |
| 2018–2019 | A(H1N1)pdm09 | 68 | 1.7398 | 1.0124 | 44 | [37, 51] | [41] |
| 2018–2019 | A(H3N2) | 47 | 1.9652 | 1.5111 | 9 | [0, 20] | [41] |
| 2019–2020 | A(H1N1)pdm09 | 47 | 1.8124 | 0.9147 | 30 | [21, 39] | [42] |
| 2019–2020 | B/Victoria | 108 | 1.5570 | 1.4553 | 45 | [37, 52] | [42] |
| 2021–2022 | A(H3N2) | 86 | 2.1416 | 1.4638 | 36 | [20, 49] | [43] |
| 2022–2023 | A(H1N1)pdm09 | 39 | 1.8935 | 1.0199 | 47 | [1, 72] | [44] |
| 2023–2024 | A(H1N1)pdm09 | 54 | 1.2748 | 0.8464 | 29 | [15, 41] | [5] |
| 2023–2024 | A(H3N2) | 13 | 1.8180 | 1.4362 | 30 | [8, 47] | [5] |
| 2023–2024 | B/Victoria | 4 | 1.0127 | 0.5581 | 74 | [65, 81] | [5] |
| 2024–2025 | A(H1N1)pdm09 | 73 | 1.6211 | 1.1130 | 37 | [24, 48] | [6] |
| 2024–2025 | A(H3N2) | 66 | 1.4044 | 1.2941 | 27 | [14, 39] | [6] |
| 2024–2025 | B/Victoria | 3 | 1.1396 | 0.7173 | 40 | [12, 59] | [6] |

#### C.3 Verification of published VE estimates and figure-derived values

Each outcome was checked against its original article before model refitting. Two of 28 assembled observations were excluded: the recorded 2017–2018 B/Victoria value of 76% was not found as a lineage-specific estimate in Rolfes et al. [4], and the recorded 2018–2019 B/Yamagata value of 34% represented combined influenza B rather than that lineage [41].

Twenty-five retained estimates were printed numbers. These include the four 2015–2016 estimates printed in Figure 1 of Jackson et al. [39]. The remaining estimate, 48% for 2017–2018 B/Yamagata, was read from the unlabelled point in Figure 2 of Rolfes et al. [4]. Applying the same reading procedure to four points with printed values recovered 38.1%, 22.0%, 62.0% and 49.9% against 38%, 22%, 62% and 50%, respectively. Omitting the figure-derived estimate left 25 observations and gave the pooled topological fit 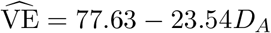 (*R*^2^ = 0.301; slope-permutation *p* = 0.0050), compared with *R*^2^ = 0.304 and *p* = 0.0033 in the primary analysis. The pooled association therefore did not depend on retaining this single figure-derived estimate.

**Table S5:** Vaccine target (seed strain) inventory. For each season and component the target is the named WHO/NIBSC/NYMC/CDC reassortant seed strain where one exists on GISAID, egg-passaged, rather than an arbitrary egg-passaged clinical isolate, because the named reassortant is the sequence used to manufacture the vaccine. Accessions and passage histories are as recorded in GISAID. The mapping from each WHO February recommendation to a target sequence was verified manually rather than programmatically.

| Vaccine seed strain | GISAID EPI_ISL | Passage | Seasons and components |
| --- | --- | --- | --- |
| A/Missouri/11/2025 (IVR-279) | EPI_ISL_20215073 | Spf3E1D6 | **2026-27 prediction**, A(H1N1)pdm09 (egg) |
| A/Darwin/1454/2025 (IVR-283) | EPI_ISL_20356066 | E2/D7 | **2026-27 prediction**, A(H3N2) (egg) |
| B/Tokyo/EIS13-175/2025 | EPI_ISL_20373027 | E3 | **2026-27 prediction**, B/Victoria (no named reassortant yet) |
| A/California/7/2009 | EPI_ISL_391380 | E2E6 | 2010-11 – 2015-16, A(H1N1)pdm09 |
| B/Brisbane/60/2008 (NIB-58) | EPI_ISL_232491 | E4 | 2010-11, 2011-12, 2013-14 – 2017-18, B/Victoria |
| A/Perth/16/2009 (NIB-64) | EPI_ISL_70443 | E4/E2 | 2010-11, 2011-12, A(H3N2) |
| A/Victoria/361/2011 (IVR-165) | EPI_ISL_168948 | E6 | 2012-13, A(H3N2) |
| B/Wisconsin/01/2010 (BX-41A) | EPI_ISL_83975 | EX | 2012-13, B/Yamagata |
| A/Texas/50/2012 (X-223A) | EPI_ISL_132571 | E5E8 | 2013-14, 2014-15, A(H3N2) |
| B/Massachusetts/2/2012 (BX-51C) | EPI_ISL_402396 | E3E7/E1 | 2013-14, 2014-15, B/Yamagata |
| B/Phuket/3073/2013 | EPI_ISL_291275 | E6 | 2015-16 – 2024-25, B/Yamagata (every season it's the vaccine strain) |
| A/Switzerland/9715293/2013 (X-247) | EPI_ISL_169603 | E4E9+1 | 2015-16, A(H3N2) |
| A/Hong Kong/4801/2014 (X-263B) | EPI_ISL_222828 | EX/E3 | 2016-17, 2017-18, A(H3N2) |
| A/Michigan/45/2015 (X-275B) | EPI_ISL_233044 | EX | 2017-18, 2018-19, A(H1N1)pdm09 |
| B/Colorado/06/2017 | EPI_ISL_276408 | E5 | 2018-19, 2019-20, B/Victoria (and combined B in 2018-19) |
| A/Singapore/INFIMH-16-0019/2016 (X-307) | EPI_ISL_293533 | E5E2E9 | 2018-19, A(H3N2) |
| A/Brisbane/02/2018 (IVR-190) | EPI_ISL_20212507 | E3/D8/E2 | 2019-20, A(H1N1)pdm09 |
| A/Victoria/2570/2019 (X-379A) | EPI_ISL_14809235 | EX | 2021-22, 2022-23, A(H1N1)pdm09 |
| A/Cambodia/e0826360/2020 (X-361A) | EPI_ISL_2378896 | EX | 2021-22, A(H3N2) |
| B/Austria/1359417/2021 (BVR-26) | EPI_ISL_5933268 | Ex | 2022-23, 2023-24, 2024-25, B/Victoria |
| A/Darwin/9/2021 (NYMC X-369A) | EPI_ISL_5522637 | EX | 2022-23, 2023-24, A(H3N2) (egg) |
| A/Victoria/4897/2022 (IVR-238) | EPI_ISL_17086692 | E3/E6 | 2023-24, 2024-25, A(H1N1)pdm09 (egg) |
| A/Thailand/8/2022 (IVR-237) | EPI_ISL_18372670 | E3/D7 | 2024-25, A(H3N2) (egg) |

Only point estimates entered the models. Confidence intervals were not available for five observations; among the 21 recorded intervals, median width was 22 percentage points (mean 26.9; range 14–71). The interval held for 2015–2016 H3N2 came from an agency archive rather than the article and is not shown as a primary-source interval in Table S4. The 2022–2023 estimate is late-season rather than a conventional full-season estimate [44]. Clinical estimation uncertainty was not propagated through the regression.

#### C.4 Word lengths, radius grids and checks of distance calculations

Both distances used HA proteins, the 20 amino acids and word lengths one to three. The protein examples of Hozumi and Wei [20] also use maximum word length three. The Betti-curve implementation used only Betti-0 (*b*_*k*_ = 0 in that paper’s Equation 16), with radii *{*1, 2, 4, …, *N}, N* being the greater of the two sequence lengths. The Grassmann implementation used 50 log-spaced radii on [1, 600], four channels and rank 10. Both grids depart from the linear schedule *r*_*i*_ = 4 ^*k*−1^*i* of the source nucleotide implementation, for the reason given in Section 3.2; the upper radius exceeds the longest protein in this dataset (585 amino acids).

**Table S6:**
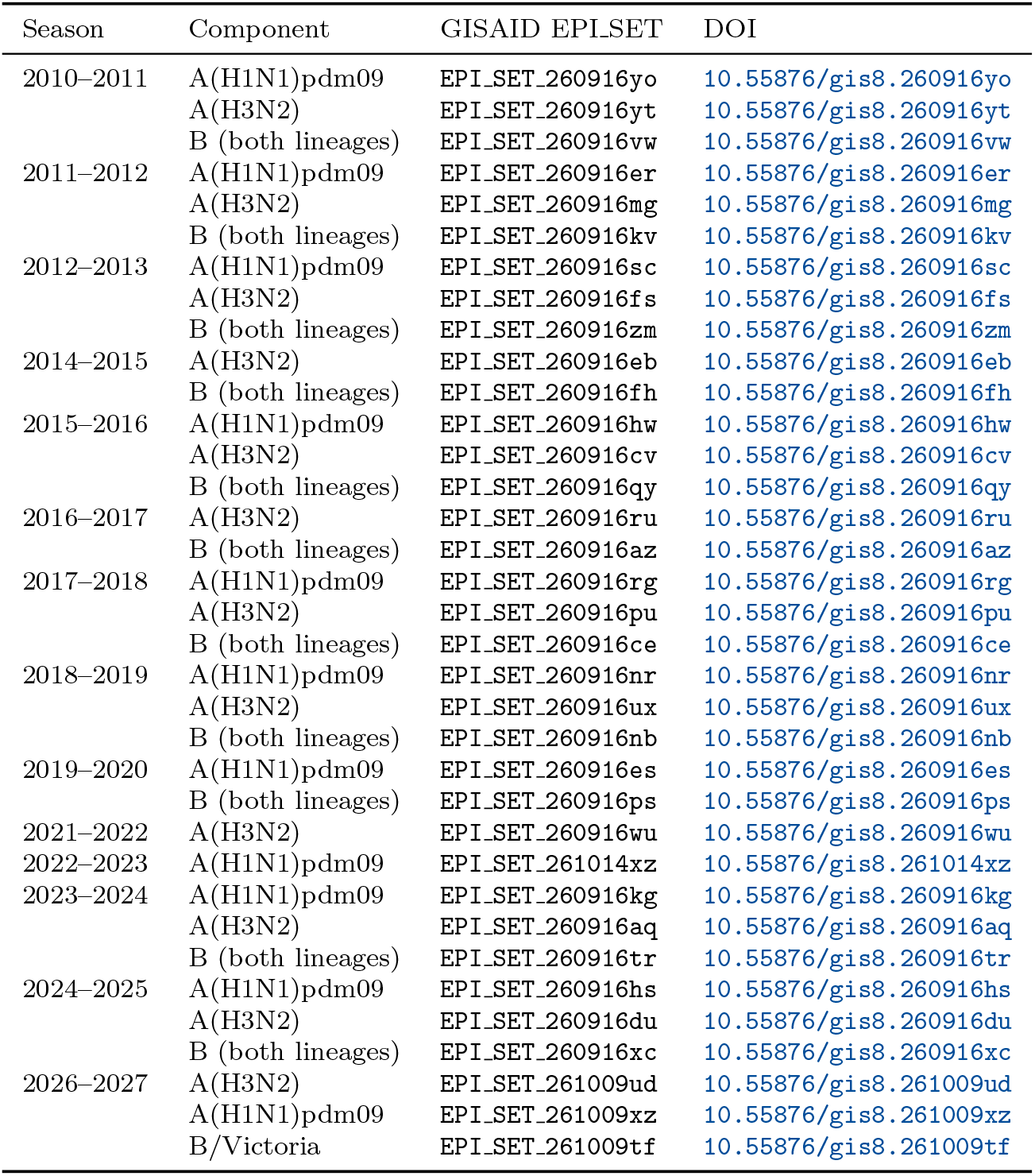
GISAID EPI SET accessions and DOIs for the preseason searches of each season contributing to the cohort or the forecast. Each DOI resolves to the acknowledgement table for that sequence set. The 2026–2027 entries are the window-closed searches of 9 October 2026 used for the frozen forecast.

The Betti-curve calculation was checked against the published formulas and internal consistency tests; it was not numerically compared with the nucleotide-only reference code. The Grassmann calculation used the reference feature, normalisation, SVD and distance routines of Wang and Wei [23], with the amino-acid word enumeration and radius grid supplied separately. With the reference nucleotide settings, feature blocks matched to machine precision across all four channels and word lengths one to three. Removing rows that are zero in every sequence preserves nonzero features and the subspace distances, provided shared coordinates and the RMS denominator are preserved.

#### C.5 Permutation tests and intervals for future VE

For the two primary pooled associations, we used two-sided slope tests with 20,000 permutations of the VE labels (NumPy random seed 0). For each permutation, we refitted the regression and compared its absolute slope with the observed absolute slope; the *p* value was the fraction of permuted slopes at least as large. This gave *p* = 0.0033 for the topological distance and *p* = 0.0009 for the geometric distance. The sequence-exclusion and printed-number-only sensitivity tests used the same procedure.

We report 95% OLS future-observation prediction intervals. These account for fitted-regression uncertainty and residual variation under the linear model, rather than uncertainty in the mean fitted VE alone.

#### C.6 Training seasons, test observations and distance scales

The evaluation retained the original common test set for comparison with the historical-mean reference. A season was eligible only if every viral group in that season had an earlier VE observation. The training set also had to contain more observations than the number of previously observed viral groups plus one, ensuring estimable residual variance under the original comparison protocol. These rules were retained when reporting only the pooled models.

The 19 test observations came from 2014–2015 (one), 2016–2017 (three), 2017–2018 (three), 2018–2019 (two), 2019–2020 (two), 2021–2022 (one), 2022–2023 (one), 2023–2024 (three) and 2024–2025 (three). The first three observations, from 2010–2011, 2011–2012 and 2012–2013, provided the initial training set. The 2015–2016 season was excluded as a whole test season because three viral groups first appeared that year; its four observations entered training for subsequent seasons. Thus, seven of the 26 observations contributed to training without being scored as test predictions.

At each test season, we refitted both the regression and distance scales using all eligible observations from earlier seasons. The Betti-curve scales were estimated from their virus–vaccine comparisons. Grassmann scales were estimated from the unique circulating and vaccine protein sequences in the same training data. These scales were then applied to both training and test distances. No sequence from the test season or a later season contributed to scale estimation.

The reference predicted the earlier mean VE for the relevant subtype or lineage. Errors were calculated once per season–component observation, with equal weight across the test observations. MAE averages absolute errors; RMSE takes the square root of the mean squared error. Coverage counts observed VE estimates inside their prediction intervals, and mean width averages the differences between interval limits. The relative MAE reduction is 100(1 −MAE_model_*/*MAE_reference_).

For Grassmann scale estimation, the RMS denominator counted all 20^*k*^ possible amino-acid words, including zero rows, so later sequences could not alter the row-count convention. A check using the original stored-row denominator gave a geometric MAE reduction of 6.2%, compared with 5.2% under the full-alphabet convention. Both calculations used the same 19 test observations; this check concerns preprocessing. Neither reduction is distinguishable from zero (Section B.2), so the convention does not change the conclusion. The training-only evaluation was implemented retrospectively. Original pooled fits and frozen forecasts retain the project-wide scales described in Section 3.2.

The split follows the seasons in which specimens and clinical outcomes were collected. It does not reconstruct exactly which sequence records or published VE estimates were publicly available at each historical forecast date. Consequently, this is a retrospective evaluation with season-ordered training and testing; prospective performance remains to be established.

#### C.7 Matching sampling windows and omitting individual pairs or seasons

The topological window comparison used 18 matched observations across eight seasons, with the same vaccine targets and scales in both windows. Scales were estimated from the preseason sequences in the combined 30,464-record collection and applied to both periods. This collection combines the original paired-window search with a later recovery of A(H1N1)pdm09 records, after removing the records flagged by the nonseasonal-virus checks. Correlations were calculated before and after subtracting group means separately within each window; uncertainty was assessed by resampling whole seasons. This comparison does not use VE labels. Enrollment-period sequences did not enter the primary VE regressions.

The component-demeaned correlation ranged from 0.755 to 0.945 after deleting one pair and from 0.783 to 0.948 after deleting one season. The correlation remained positive and high in each deletion, so no single pair or season was solely responsible for the overall relationship. Within-group correlations were 0.98 for H1N1 (*n* = 5), 0.93 for B/Victoria (*n* = 4), 0.70 for H3N2 (*n* = 6) and −0.30 for B/Yamagata (*n* = 3). In 2018–2019 H3N2, the preseason distance was within 0.06 of its group mean, while the enrollment-period distance was 0.46 above its group mean. Thus, high overall correlation did not imply agreement in every group or season.

The enrollment-period collection contains 29,461 sequences. No GISAID EPI SET identifier was registered when these searches were performed, so these sequences are not covered by the identifiers in Supplementary Table S6. Their accessions are instead listed by season, viral group and sampling window in `metric_validation/paired_window_accessions.cs` in the repository, from which the corresponding GISAID records and their contributor acknowledgements can be retrieved. These searches nonetheless remain less fully documented than the preseason datasets. The matched seasons were selected by data availability rather than random sampling.

## Notes

### Competing Interest Statement

The authors have declared no competing interest.

